# Pulmonary vein flow split effects in patient-specific simulations of left atrial flow

**DOI:** 10.1101/2023.03.24.534045

**Authors:** E. Durán, M. García-Villalba, P. Martinez-Legazpi, A. Gonzalo, E. McVeigh, A.M. Kahn, J. Bermejo, O. Flores, J.C. del Álamo

**Affiliations:** Department of Mechanical, Thermal and Fluids Engineering, Universidad de Málaga, Málaga, Spain; Department of Aerospace Engineering, University Carlos III of Madrid, Leganés, Spain; Institute of Fluid Mechanics and Heat Transfer, TU Wien, 1060 Vienna, Austria; Department of Mathematical Physics and Fluids, National University of Distance Education (UNED), Madrid, Spain; Department of Mechanical Engineering, University of Washington, Seattle, WA, United States; Department of Bioengineering, University of California San Diego, La Jolla, CA, United States; Department of Radiology, University of California San Diego, La Jolla, CA, United States; Division of Cardiovascular Medicine, University of California San Diego, La Jolla, CA, United States; Gregorio Marañón University Hospital, Madrid, Spain; Spanish Cardiovascular Network (CIBERCV), Carlos III Health Institute, Spain; Faculty of Medicine, Complutense Unversity, Madrid, Spain; Gregorio Marañón Health Research Institute (IISGM), Madrid, Spain; Center for Cardiovascular Biology, University of Washington, Seattle, WA, United States; Institute for Stem Cell and Regenerative Medicine, University of Washington, Seattle, WA, United States

**Keywords:** computational fluid dynamics, cardiovascular flows, imaged-based simulation, left atrium, thrombosis, pulmonary veins

## Abstract

Disruptions to left atrial (LA) blood flow, such as those caused by atrial fibrillation (AF), can lead to thrombosis in the left atrial appendage (LAA) and an increased risk of systemic embolism. LA hemodynamics are influenced by various factors, including LA anatomy and function, and pulmonary vein (PV) inflow conditions. In particular, the PV flow split can vary significantly among and within patients depending on multiple factors. In this study, we investigated how changes in PV flow split affect LA flow transport, focusing on blood stasis in the LAA, using a high-fidelity patient-specific computational fluid dynamics (CFD) model. We analyzed LA anatomies from eight patients with varying atrial function, including three with AF and either a LAA thrombus or a history of TIAs. Using four different flow splits (60/40% and 55/45% through right and left PVs, even flow rate, and same velocity through each PV), we found that flow patterns are sensitive to PV flow split variations, particularly in planes parallel to the mitral valve. Changes in PV flow split also had a significant impact on blood stasis and could contribute to increased risk for thrombosis inside the LAA, particularly in patients with AF and previous LAA thrombus or a history of TIAs. Our study highlights the importance of considering patient-specific PV flow split variations when assessing LA hemodynamics and identifying patients at increased risk for thrombosis and stroke.

## 1 Introduction

Cardiovascular diseases are the leading cause of mortality worldwide. Atrial fibrillation (AF) affects around 35 million people worldwide [1], and is the most common arrhythmia. It is estimated that 20-25% of ischemic strokes, affecting over 18 million people each year, are caused by thrombi generated in the left atrium (LA) in patients with AF. Furthermore, up to an additional 30% of all ischemic strokes are suspected to be atriogenic in patients with subclinical AF or normal cardiac rhythm [2]. Inside the LA, most of these thrombi are formed in the left atrial appendage (LAA), a cavity-shaped protuberance whose morphology and hemodynamics vary significantly among patients [3]. However, current medical procedures to estimate ischemic stroke risk are based on demographic and clinical factors and do not consider patient-specific information about LA hemodynamics, a crucial causal thrombosis factor [4, 5].

Computational fluid dynamics (CFD) analysis based on patient-specific medical images is a powerful tool for investigating LA thrombosis. In the past decade, advances in CFD algorithms, numerical algorithms and computational resources have supported increasingly complex cardiac flow simulation studies. These works cover four-chamber simulations of the whole heart [6], two-chamber simulations of the left heart including the atrium and ventricle [7–10], and single-chamber simulations of the LA or LV. These single-chamber simulations have focused mostly on LV hemodynamics [11–16], although there is a growing body of literature exploring different aspects of LA hemodynamics. One of the most studied aspects is the effect of LA wall motion on the flow and LAA stasis [17–21], motivated by the fact that AF causes weak irregular LA wall motion and LAA stasis is associated with increased thrombosis risk [1, 22–25]. Motivated by the clinical association between LAA morphology and stroke risk in AF patients [26–28], several groups have used CFD to investigate how LAA shape influences surrogate hemodynamic metrics of thrombosis risk [29–32]. In parallel, recent work [10, 33–38] is introducing advanced CFD, fluid-structure interaction, and multi-physics models, and integrating them with multi-modality imaging to realistically simulate the electrophysiological and biomechanical mechanisms underlying LAA stasis and thrombosis.

The aforementioned simulation studies have provided useful guidance for setting up models and boundary conditions that reproduce each patient’s hemodynamics representative of a specific time or physiological state. However, both errors in model approximation and the natural variation of physiological state during a patient’s daily life (e.g., rest vs. exercise) constitute a source of uncertainty. These errors must be understood, evaluated, and accounted for before model predictions of LAA thrombosis based on hemodynamic computations can be used to support clinical decisions. The PV flow split, i.e., how the flowrate of blood entering the LA from the lungs is divided between the left and right PVs, is a particularly challenging parameter affecting the inflow boundary conditions. Its quantification via transthoracic Doppler echocardiography (TTE) is complicated by the difficulty of imaging all (usually four) pulmonary veins [39] and, while it can be accessed by other modalities, such as 4D flow MRI, these techniques are more cumbersome and time-consuming than TTE. Consequently, the PV flow split is typically assumed to be even for simplicity, even if the difference in size between the left and right lungs makes this split be closer to 47%/53% (left/right) [40], which can vary widely from patient to patient (Cheng et al.[40] report a range of 43-48/57-52% for N=206 patients). Moreover, the PV flow split can experience significant intra-subject variability due to interactions between gravity and body position, with reported differences between lying on the left and right sides of the body being 59/41% and 37/63%, respectively [41]. These significant inter- and intra-patient variabilities in PV flow split warrant investigating its effect on LA hemodynamics and LAA blood stasis.

There are few works investigating the influence of PV inflow in CFD simulations of atrial blood flow, and most of them do not directly or systematically address the left/right flow split. Lantz et al.[42] carried out a systematic parametric study, including a thorough sweep over a wide range of different PV flow splits and validation vs. in vivo 4D flow MRI measurements. Their results demonstrated that atrial flow can be sensitive to the PV flow split, especially for large departures from normal values. However, they only considered N=3 patients and did not address LAA flow or residence time. Previously, García-Isla et al.[29] had investigated the effect of PV inflow as part of a broader sensitivity analysis of the effect of LA and LAA morphological features on atrial flow dynamics. By varying PV cross sectional area while keeping inflow velocity constant and an even 50/50 split, these authors effectively evaluated the effect of total PV flow rate on LA hemodynamics, showing significant effects. Subsequently, Mill et al.[25] conducted patient-specific simulations on a large cohort of N=52 patients, concluding that the position and orientation of the right PVs can significantly influence LA hemodynamics. Nevertheless, neither García-Isla et al.[29] or Mill et al.[25] addressed the effect of the PV flow split. More recently, Otani et al. [43] and Yi et al. [44] have studied LA hemodynamics before and after surgical removal of left upper lung lobe, concluding that the resulting left superior pulmonary vein recession disturbs blood flow all throughout the LA. These two studies represent an additional application of CFD for patient-specific prediction of atrial flow in situations where the PV flow split changes dramatically.

This manuscript presents a CFD investigation of PV flow split effects on subjects with normal and impaired, pro-thrombotic atria. Patient-specific anatomical models are built based on 4D computed tomography (CT) images, including total flow rate through the PVs. For each model, we consider four PV flow split variations encompassing the even split commonly used in CFD analysis and more physiological, asymmetric splits with higher flow through the right PVs. Particular attention is given to PV-flow split induced variability in LAA blood stasis.

## 2 Methodology

### 2.1 Imaging and generation of patient-specific anatomical models

We studied N = 8 cases through 3D, time-resolved (a.k.a. 4D), patient-specific segmentations of human left atria obtained from computed tomography (CT) images. Six of these segmentations are the same as in [24] and [37]. Two additional subjects were imaged at Hospital General Universitario Gregorio Marañón (HGUGM), Madrid, Spain, following standard clinical protocols using a Philips Brilliance 64 scanner. The doses and injection rates of contrast were chosen based on the patient’s weight using standard clinical protocols. The images were reconstructed using CT scanner manufacturers’ algorithms, yielding DICOM files with pixel dimensions between 0.375mm and 0.42mm in the x-y plane and 0.45mm in the z-direction. For the other six cases, pixel dimension was 0.32mm to 0.48mm in the x-y plane and 0.5mm to 1mm in the z-direction. Time-resolved images were obtained at regularly spaced instants across the cardiac cycle, ranging between 5% and 10% of the R-R interval.

The computational LA meshes were generated in four steps using ITK-SNAP [45] and custom-written scripts in MATLAB. The first step comprised segmenting the 3D LA anatomy from CT images and identifying the PV inlets, mitral annulus, and LAA. For each 3D segmentation, a triangular surface mesh was created and then resampled to match the computational fluid dynamics (CFD) solver’s resolution [46]. The resulting triangular meshes were registered across the cardiac cycle and their positions, yielding a coherent triangle vertex and centroid cloud [47, 48]. Finally, the positions of these points were expressed as Fourier temporal series to provide interpolated boundary conditions to the CFD solver at time points not contained in the 4D CT sequence. Table 1 summarizes the main anatomical and functional features of the study subjects’ atria. Figure 1 shows the N=8 patient specific anatomies at the beginning of the R-R interval. Additional details on image acquisition and reconstruction and mesh generation can be found elsewhere [24].

**Table 1.** Anatomical and functional parameters of the LA and the LAA. The mean volume values represent time-averaged volumes. Ostium AR: ratio between the length of the ostium major and minor axes. Ostium AR and ostium area are time-averaged values over the cardiac cycle. EF: ejection fraction. Global (LA) EF: (VLA,max-VLA,min)/VLA,max. LAA EF: (VLAA,max-VLAA,min)/VLAA,max. Expansion index: (VLA,max-VLA,min)/VLA,min. Passive EF: (VLA,max-VLA,preA)/VLA,max. Booster EF: (VLA,preA-VLA,min)/VLA,preA. Conduit Volume / LV Stroke Volume: 1 - (VLA,max-VLA,min)/LVSV. PVs: blood volume that enters the LA during LV systole. PVd: blood volume that enters the LA during LV diastole. PVa: blood volume that exits the LA due to reverse flow volume through the PV during atrial contraction. E/A ratio: ratio of peak mitral velocities during early diastole (E-wave) and atrial contraction (A-wave). *Re_MV_*: Reynolds number at the MV, computed as *Re_MV_* = *v_MV_ D_MV_*/*ν*, using a characteristic diameter *D_MV_* = (4*A_MV_* /π)^1/2^, where *A_MV_* is the area of the MV, the peak velocity during the E-wave *u_MV_* = *Q_MV_* / *A_MV_*, and the blood viscosity, *ν*.

| Subject | 1 | 2 | 3 | 4 | 5 | 6 | 7 | 8 |
| --- | --- | --- | --- | --- | --- | --- | --- | --- |
| LAA thrombus (or history of TIAs) | No | No | No | No | No | TIAs | Yes | Yes |
| Atrial fibrillation | No | No | No | No | No | Persistent | Persistent | Persistent |
| Sinus rhythm | Yes | Yes | Yes | Yes | Yes | No | No | No |
| eqV flow split (L/R) (%) | 42/58 | 46/54 | 42/58 | 40/60 | 34/64 | 51/49 | 39/61 | 46/54 |
| Mean LA volume (ml) | 86.6 | 70.1 | 115 | 84.2 | 96.1 | 145 | 157 | 180 |
| Min. LA Vol. (VLA,min, ml) | 59.6 | 49.0 | 87.2 | 60.3 | 68.6 | 119 | 150 | 157 |
| Max. LA Vol. (VLA,max, ml) | 108 | 91.2 | 145 | 108 | 121 | 155 | 165 | 205 |
| Mean LAA volume (ml) | 6.94 | 4.85 | 14.3 | 3.13 | 3.64 | 10.7 | 15.5 | 22.0 |
| Min. LAA Vol. (VLAA,min, ml) | 4.32 | 3.14 | 10.2 | 1.45 | 2.24 | 9.10 | 13.8 | 19.8 |
| Max. LAA Vol. (VLAA,max, ml) | 8.97 | 6.28 | 17.9 | 4.23 | 4.65 | 11.6 | 17.4 | 24.7 |
| LV stroke volume (LVSv, ml) | 72.2 | 84.4 | 98 | 106 | 102 | 66.7 | 75 | 130 |
| Ostium area (mm<sup>2</sup>) | 4.17 | 4.01 | 7.65 | 2.26 | 3.00 | 7.22 | 7.29 | 8.42 |
| Ostium AR | 1.53 | 1.39 | 1.49 | 1.10 | 1.19 | 1.45 | 1.45 | 1.83 |
| Pre-A-wave Vol. (VLA,preA, ml) | 94.3 | 72.3 | 119 | 87.3 | 98.5 | 149 | 153 | 178 |
| LA sphericity | 0.80 | 0.82 | 0.78 | 0.77 | 0.83 | 0.86 | 0.83 | 0.81 |
| Global (LA) EF | 0.45 | 0.46 | 0.40 | 0.44 | 0.43 | 0.23 | 0.09 | 0.23 |
| LAA EF | 0.52 | 0.50 | 0.43 | 0.66 | 0.52 | 0.22 | 0.21 | 0.20 |
| Expansion index | 0.81 | 0.86 | 0.66 | 0.80 | 0.76 | 0.30 | 0.10 | 0.31 |
| Passive EF | 0.13 | 0.21 | 0.18 | 0.19 | 0.18 | 0.04 | 0.07 | 0.13 |
| Booster EF | 0.37 | 0.32 | 0.27 | 0.31 | 0.30 | 0.20 | 0.02 | 0.12 |
| Conduit Volume / LV Stroke Volume | 0.33 | 0.50 | 0.41 | 0.55 | 0.49 | 0.46 | 0.80 | 0.63 |
| PVs (ml) | 43 | 38 | 51 | 48 | 53 | 36 | 17 | 49 |
| PVd (ml) | 65 | 58 | 60 | 65 | 54 | 52 | 63 | 94 |
| PVs-PVd ratio | 0.66 | 0.66 | 0.84 | 0.72 | 0.99 | 0.69 | 0.27 | 0.52 |
| PVa (ml) | 17 | 7.3 | 12 | 2.7 | 2.73 | 13 | 0.14 | 1.4 |
| PVa duration (cycles) | 0.11 | 0.11 | 0.13 | 0.08 | 0.19 | 0.12 | 0.039 | 0.068 |
| A-wave duration (cycles) | 0.22 | 0.25 | 0.30 | 0.24 | 0.28 | 0.18 | 0.21 | 0.23 |
| PVa-A-wave duration ratio | 0.49 | 0.43 | 0.44 | 0.33 | 0.68 | 0.64 | 0.18 | 0.30 |
| E/A ratio | 1.64 | 2.36 | 3.03 | 2.24 | 2.23 | 2.05 | 3.86 | 3.65 |
| $Re_{MV}$ | 4297 | 6266 | 6315 | 8402 | 6244 | 4361 | 4111 | 8058 |

**Fig. 1.**
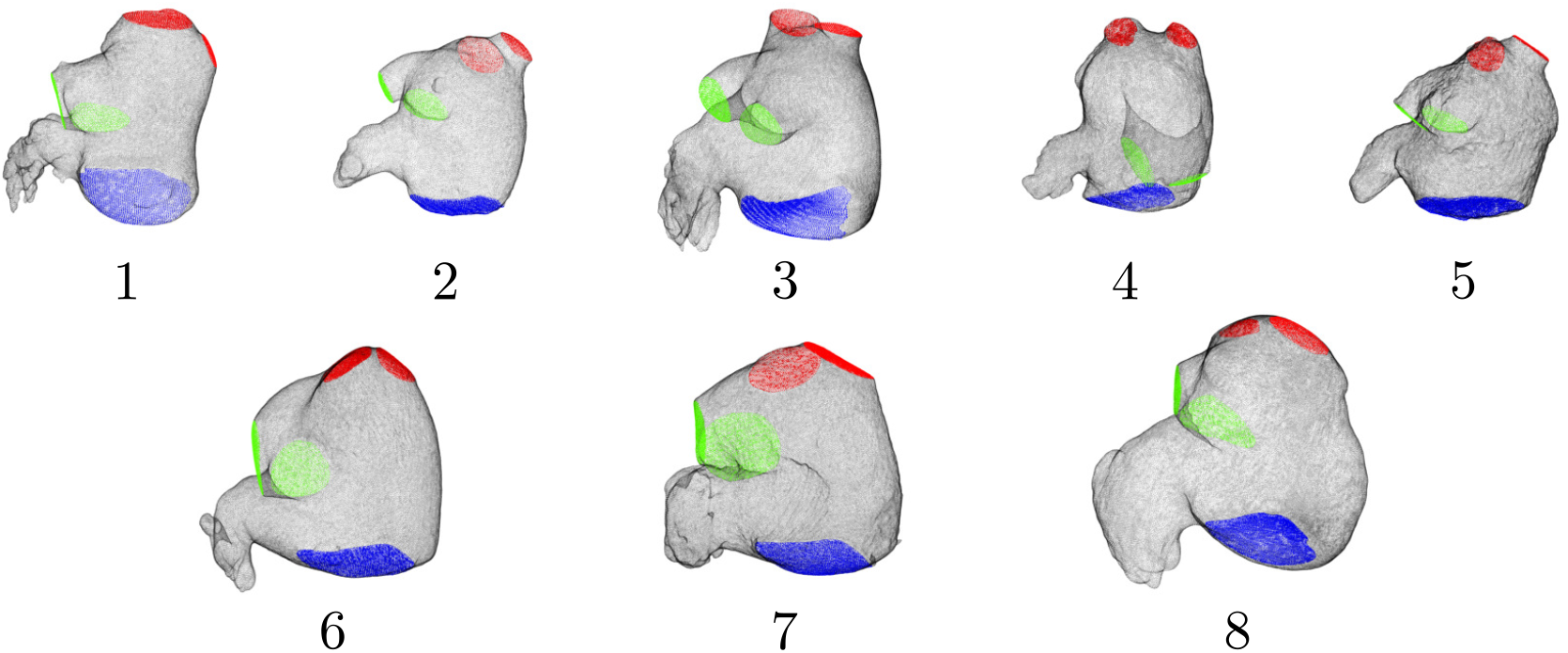
Anatomical geometry of patient-specific left atrium subjects. 3D mesh extracted from Computerized Tomography (CT) images of the LA walls and PVs inlets (left PVs in green, right PVs in red) and MV outlet surfaces (blue). The images correspond to an instant at the beginning of the R-R interval.

### 2.2 Computational Fluid Dynamics Analysis

The CFD simulations were performed with the in-house code TUCAN, as described in [24]. Here, we summarize the main features of these simulations, emphasizing the aspects that are particularly relevant to the present study of PV flow split variability.

TUCAN solves the Navier-Stokes equations for the incompressible flow using second-order centered finite differences and a three-stage, semi-implicit, low-storage Runge-Kutta scheme. In the present study, we considered a Newtonian fluid of constant kinematic viscosity *ν* = 0.04cm^2^/s. Each simulation was run in a uniform grid of 256^3^ points with *x* = 0.051cm in all directions and with a constant time step of *t* = 5 ·10^-5^s. This time step is chosen to ensure that the Courant-Friedrich-Lewis number is always *CFL* < 0.3. The numerical error of the time integrator is proportional to a power of the *CFL* number [49], so it must be small to ensure numerical stability and accuracy in the results. The simulations were initialized from a static flow field and, before running in the nominal conditions, some cardiac cycles were run with a lower resolution of *x* = 0.09cm in all directions and a constant time step of *t* = 10^-4^s. In total, we run 10 heartbeats at low resolution to accelerate convergence to periodic flow, followed by 5 more heartbeats at the nominal spatial and time resolution.

The flow in the LA was driven by each subject’s left heart wall motion, obtained from patient-specific 4D CT images as described above. These images were used to measure the LA and LV volumes along the cardiac cycle and to compute patient-specific flow rates through the mitral valve and pulmonary veins. The patientspecific LA wall motion was prescribed as no-slip boundary conditions using the immersed boundary method (IBM). Similar to [24], the mitral valve was modeled as a flat impermeable surface during LV systole and as an open boundary during LV diastole.

### 2.3 Pulmonary vein boundary conditions

To enforce flow rates through the PVs, a buffer region was defined upstream of each PV inlet plane, and the IBM was applied in this region to bring the flow velocity to the desired vector value. The target inlet velocity at each PV inlet plane was defined as 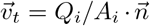, where *A_i_* and 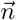 are the instantaneous cross-sectional area and the vector normal to each inlet’s plane, respectively, and *Q_i_* (*i* = 1,…, 4) are the instantaneous flow rates through each PV.

The total flow entering the LA through the four pulmonary veins, Q_PV_ (*t*), was determined from mass conservation (figure 2):

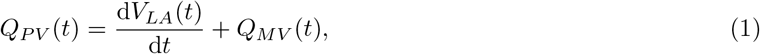

where *V_LA_*(*t*) is the time-dependent LA volume obtained from each patient’s 4D CT scan and *Q_MV_* (*t*) is the flow rate exiting the LA through the mitral valve, calculated from the time rate of change of the LV volume assuming no mitral regurgitation.

**Fig. 2.**
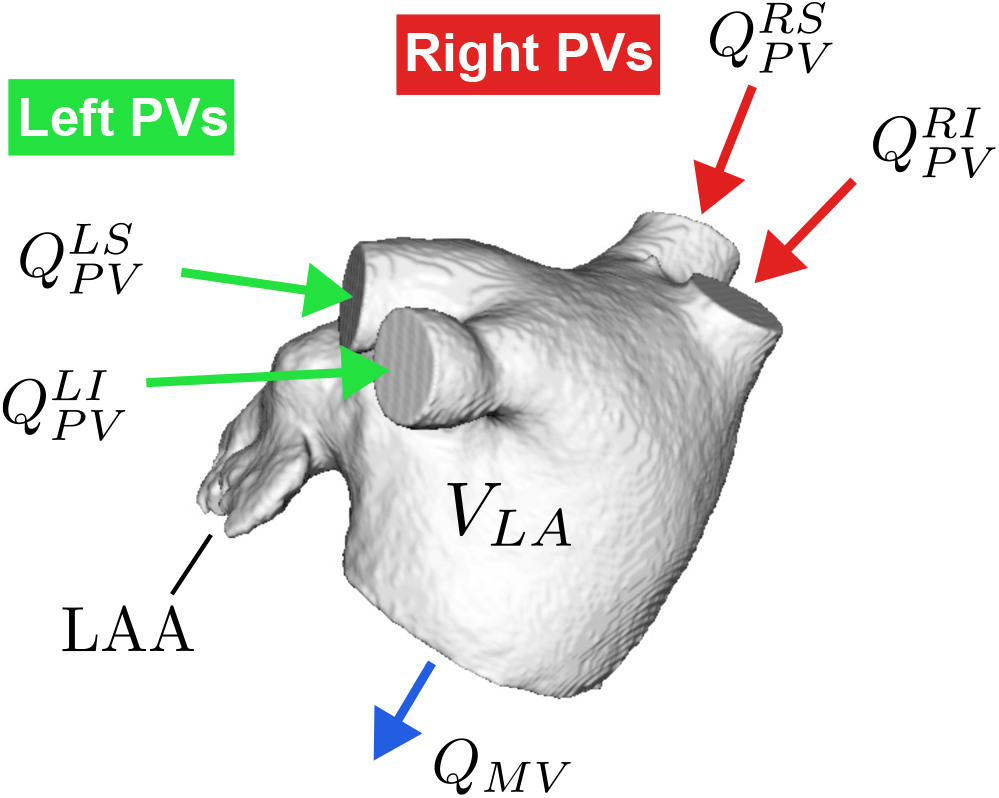
Geometry and flow distribution in the left atrium. *Q_PV_* is the inflow through the pulmonary veins (RS Right Superior, RI Right Inferior, LS Left Superior, LI Left Inferior). *V_LA_* is the time-dependent volume of the left atrium. *Q_MV_* is the outflow through the mitral valve. LAA is the left atrial appendage.

To analyze how the PV flow split affects atrial hemodynamics and blood stasis, we considered four flow split scenarios:

a. **R60**: In this scenario, 60% (40%) of the flow enters the LA through the right (left) PVs,

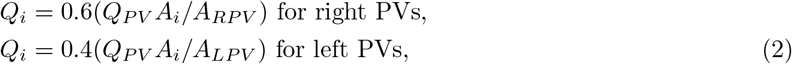

where *A_RPV_* and *A_LPV_* are the sum of the areas of the right and left PVs respectively. Usually, both the left and right PVs have two inlets into the LA body (figure 2). The same velocity value is imposed at each side’s inlet.
b. **R55**: This scenario is similar to the R60 but with a 55/45% R/L flow split.
c. **R50\***: The flow rate is split evenly among the four inlets formed by the right and left PVs,

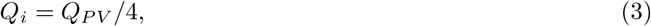

so each inlet’s velocity depends on its sectional area.
d. **eqV**: Same velocity through each PV,

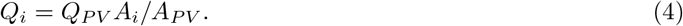 Where *A_PV_* is the sum of the areas of all PVs. In this scenario, the flow rate through each PV is not prescribed as it depends on the corresponding sectional area. The eqV flow splits for the 8 cases are shown in Table 1.

### 2.4 Flow analysis

This section describes the post-processing of CFD-computed blood velocity fields. First, we describe a framework to divide the LA in different sub-domains to study the regional impact of PV flow split variations. Then, we define correlation factors that quantify the similarity of LA hemodynamics for different PV flow splits. Finally, we describe the computation of residence time to quantify blood stasis.

To analyze the flow inside the different regions of the LA (see [50], for reference), we divided each anatomical model into three domains (figure 3A): the posterior region receiving the PVs (denoted LAPV), the appendage (denoted LAA), and the remaining part of the LA body that includes the septal portion and the vestibule (denoted LAb). While the LAA has a relatively well-defined opening, the other parts do not have clear anatomical demarcations. Thus, we devised an objective procedure to separate the LAPV and LAb domains. We defined an intrinsic coordinate ξ that defines planes parallel to the mean MV crossing the LA body (figure 3B). Instantaneous MV planes are computed by least-squares fitting the points that define the MV in each subject’s anatomical model. The coordinate ξ is normalized to range from ξ = 0 at the centroid of the PV inlets to ξ = 1 at the centroid of the MV at each instant of time. This coordinate allows for establishing an objective criterion to dynamically demarcate the LAPV and LAb (figure 3B). The separation from the LA body and the LAA is defined by a plane that crosses the LAA ostium as part of the CT image segmentation.

**Fig. 3.**
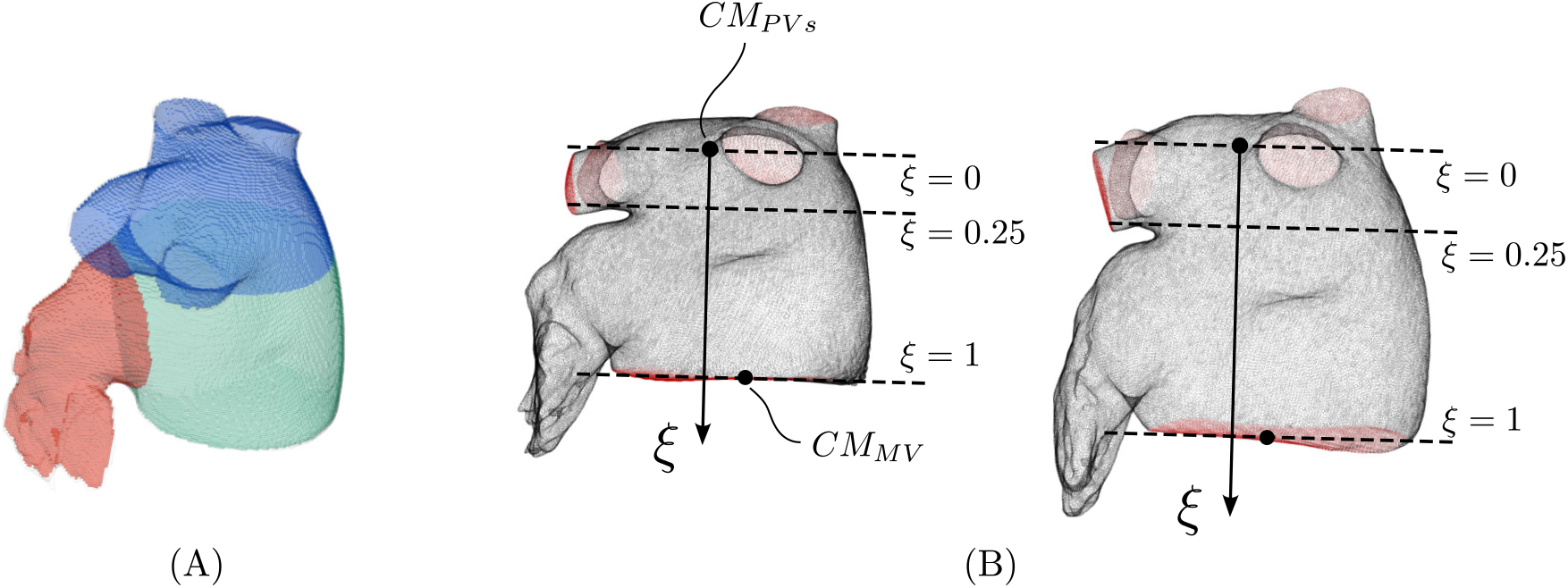
(A) Division of the left atrium geometry into three regions: pulmonary veins and the portion of the left atrium adjacent to their ostia (blue), main left atrial body (green), and left atrial appendage (red). (B) Definition of parallel planes (dashed line) dividing the left atrium. Coordinate ξ = 0 is defined with the plane that crosses the centroid of the ensemble of pulmonary veins inlets (*CM_PVs_*) at a given instant of the cardiac cycle, and coordinate ξ = 1 with the plane that crosses the centroid of the mitral valve surface (*CM_MV_*).

To quantify the dependence of LA hemodynamics on the PV flow split, we define correlation factors for the PV inflows and the velocity fields in the LA. The correlation factor of PV flow rates is defined as

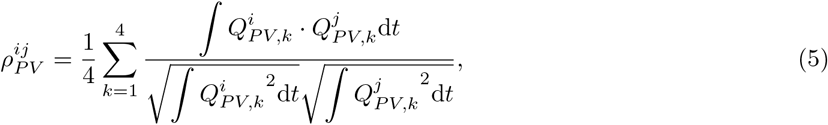

where 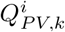 is the flowrate waveform corresponding to the *k*-th PV with the flow split condition indicated by *i*=[R60,R55,R50*,eqV]. Likewise, the correlation factor of the 4D velocity field is defined as

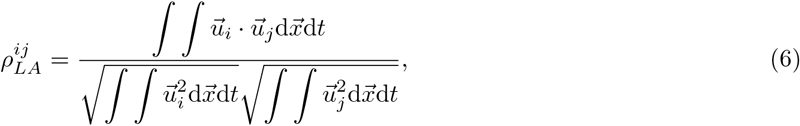

where 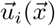 is the velocity field at point 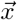 of the LA at the flow split condition indicated by *i*.

To measure blood stasis, we define the residence time *T_R_* as the time spent by blood particles inside the LA chamber. This quantity was calculated together with the velocity field by solving the forced transport equation [51],

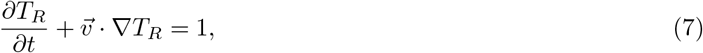

where *T_R_* is set to zero at the start of the simulation and at the PV inlets. The spatial discretization of this equation employed a third-order WENO scheme [52] to ensure accuracy while avoiding spurious fluctuations near steep gradients. We note that, up to numerical errors, this procedure is equivalent to tracking Lagrangian flow tracers and measuring the time elapsed along each tracer’s trajectories [51].

## 3 Results

### 3.1 Study population

The LA patient-specific anatomical models and functional characteristics of the study group are indicated in figure 1 and table 1. The cohort’s median age was 65 (range 48-92), and four (50%) were female. Subjects 4 and 5 were imaged at HGUGM, whereas the rest of the patients were the same reported in [24]. Subjects 1 – 5 were imaged in sinus rhythm and had no LAA thrombus. These subjects had normal LA function, as can be deduced from chamber volumes and PV flows derived from CT images. Subjects 6 and 7 were imaged in atrial fibrillation and had an LAA thrombus, which was digitally removed before the segmentation of the LA geometry. Subject 6 was imaged in atrial fibrillation and did not have an LAA thrombus; however, this patient had a previous history of transient ischemic attacks (TIAs). Subjects 6 – 8 had an enlarged LA with impaired global function. While subject 6 had decreased reservoir function but relatively normal booster function, subjects 7 and 8 had impaired reservoir and booster function. In particular, subject 7 had severely impaired function, operating primarily as a conduit between the PVs and the LV. Based on this information, we categorized subjects 1 – 5 as LAA-thrombus/TIA negative (LAAT/TIA-neg), and subjects 6 – 8 as LAAT/TIA-pos.

### 3.2 Effect of PV flow split on left atrial flow patterns

Figure 4 shows 3D velocity vector maps representative of normal atrial function (subject 2) for the different flow split scenarios considered in this study. Flow maps in a MV-parallel plane of the LA body and the LAA are represented at LA diastole and LV early filling, together with total PV flow rate waveforms. During atrial diastole, the mitral valve is closed while the left atrium and its appendage are filled with blood entering through the PVs. In this phase of the cardiac cycle, the flow patterns in the MV-parallel plane exhibit appreciable differences between flow splits, especially in the LA body (figure 4A). These differences are noteworthy considering that the PV inflow waveforms do not experience a striking dependence on the PV flow split (figure 4C). In this normal subject, the LAA flow patterns only exhibit modest differences when the PV flow split is varied. During LV relaxation, some flow is still entering through the PVs, however, the MV is now open and LV suction accelerates the blood towards the mitral annulus. In this phase, the flow organization in both the LA body and the LAA seems to be less sensitive to the PV flow split, probably under the influence of the strong atrial emptying jet (figure 4B). The 3D velocity vector maps indicative of impaired, pro-thrombotic atria (subject 6) are represented in figure 5. Similar to the normal case, the PV flow split has a noticeable effect on the flow patterns contained in the MV-parallel plane. In addition, the LAA flow patterns appear to be more sensitive to the PV flow split than in the normal case. Below, we evaluate these qualitative observations more quantitatively.

**Fig. 4.**
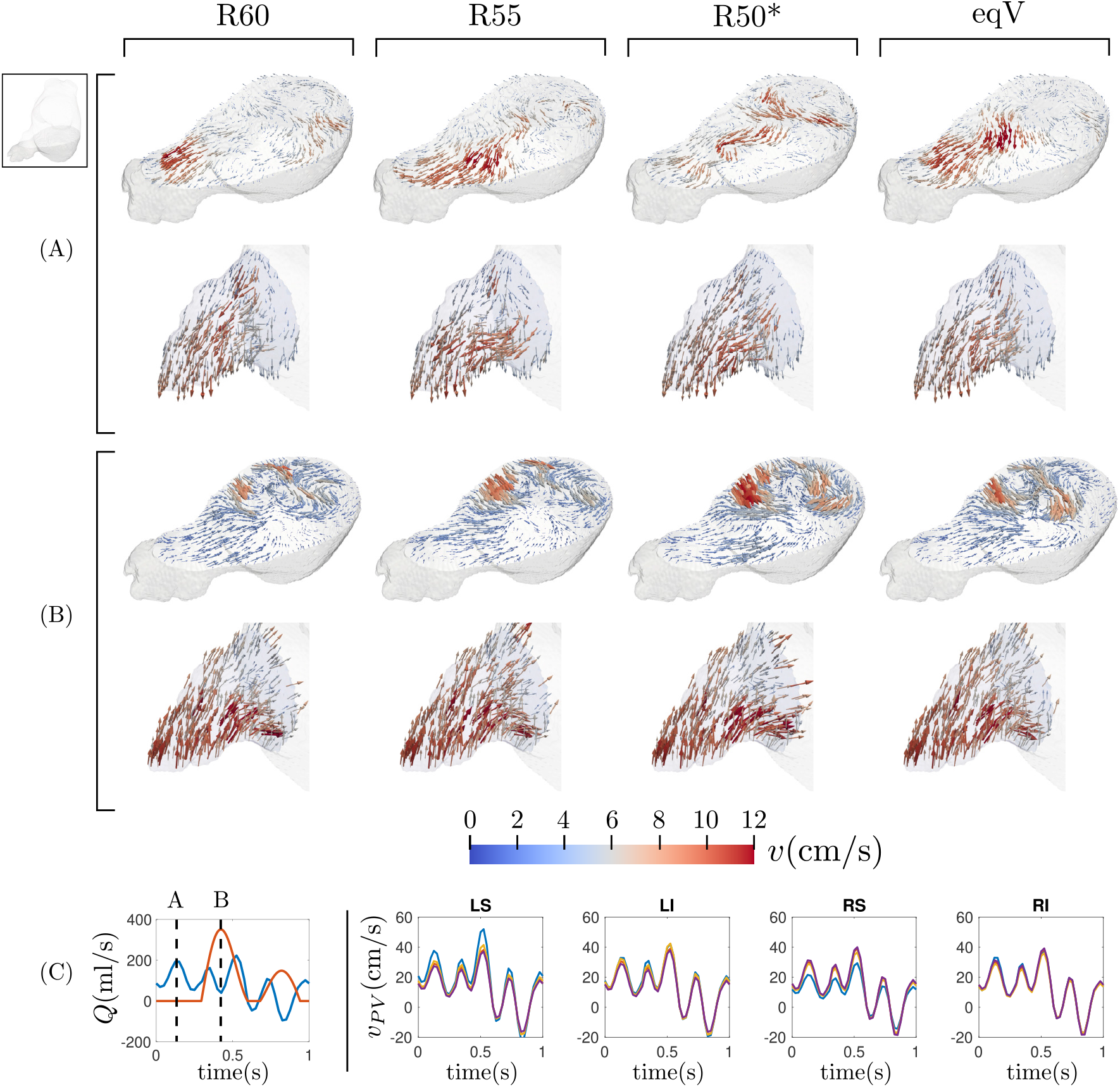
Visualization of the flow in the left atrium for the different flow split conditions during the atrial diastole (A) and during the ventricle diastole (B) in subject 2. For both instants, we represent the flow contained in a plane that crosses the atrial body and the flow inside the left atrial appendage. Panel (C) shows the temporal evolution of the flow rate through the mitral valve (red) and the pulmonary veins (blue), and indicates the position of the instants of (A) and (B) in the cardiac cycle. The right part of panel (C) shows the temporal evolution of the mean velocity through each pulmonary vein inlet for the different flow splits (R60, violet; R55, yellow; R50*, blue; eqV, red).

**Fig. 5.**
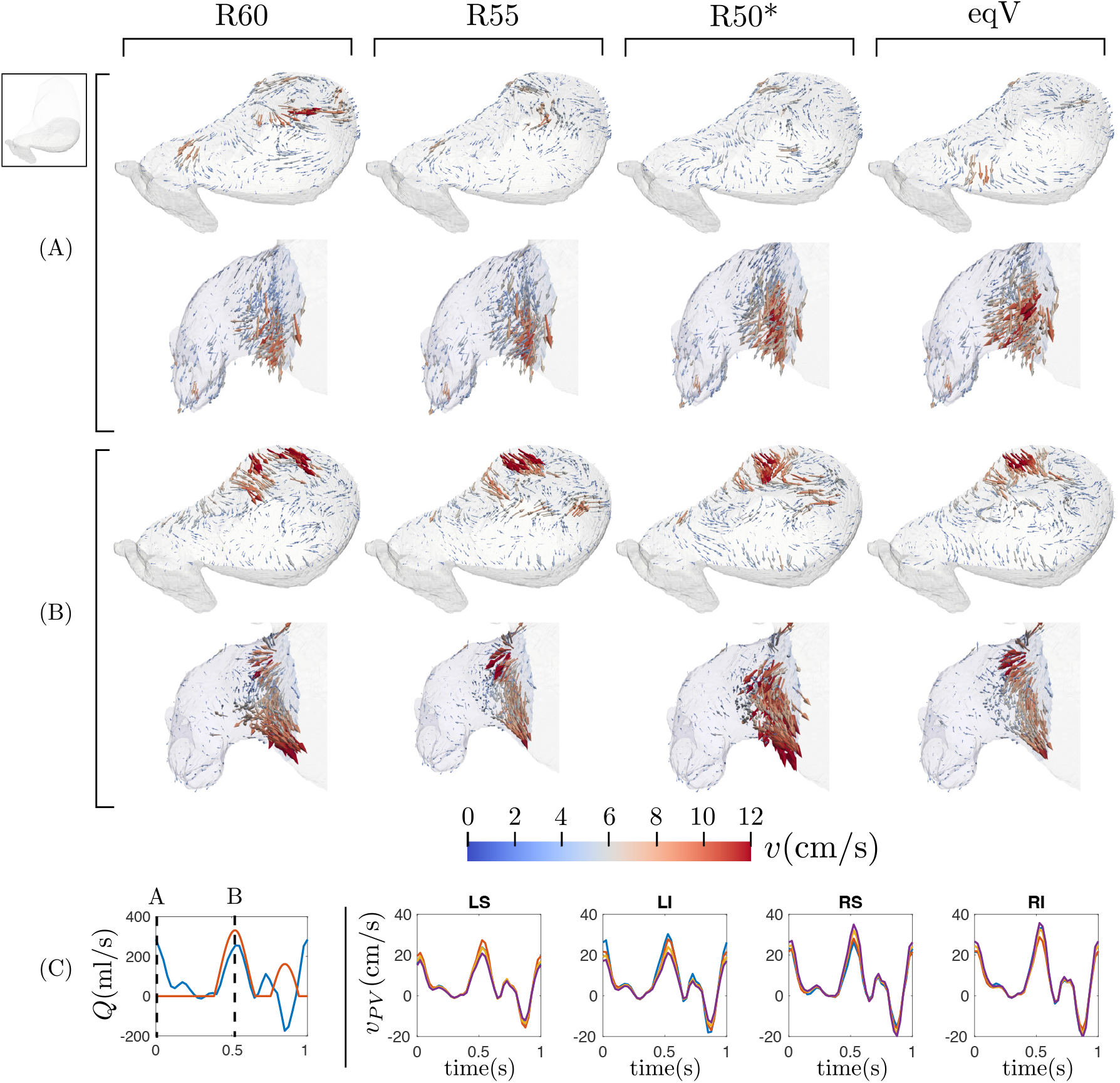
Visualization of the flow in the left atrium for the different flow split conditions during atrial diastole (A) and ventricular diastole (B) in subject 6. For both instants, we represent the flow contained in a plane that crosses the atrial body and the flow inside the left atrial appendage. Panel (C) shows the temporal evolution of the flow rate through the mitral valve (red) and the pulmonary veins (blue), and indicates the position of the instants of (A) and (B) in the cardiac cycle. The right part of panel (C) shows the temporal evolution of the mean velocity through each pulmonary vein inlet for the different flow splits (R60, violet; R55, yellow; R50*, blue; eqV, red).

Figures 6 and 7 show the blood kinetic energy (*KE*) and residence time (*T_R_*) in a plane section of the LA body and the LAA for the same two subjects of figures 4 and 5. During atrial diastole, the *KE* reaches moderate values at the PV inflow jets and is significantly lower elsewhere. During LV relaxation, the PV region still presents moderate *KE* values while its peak values are associated with the emptying jet. This overall pattern is conserved for the four flow-split scenarios considered; however, the *KE* maps show appreciable differences when comparing different flow splits and subjects. These differences are obvious in areas of high *KE* values corresponding to inflow and outflow jets. However, there are also important relative differences in the flow *KE* inside the LAA, as evidenced when representing this quantity using a ~10-fold lower range (figures 6C, 7C).

**Fig. 6.**
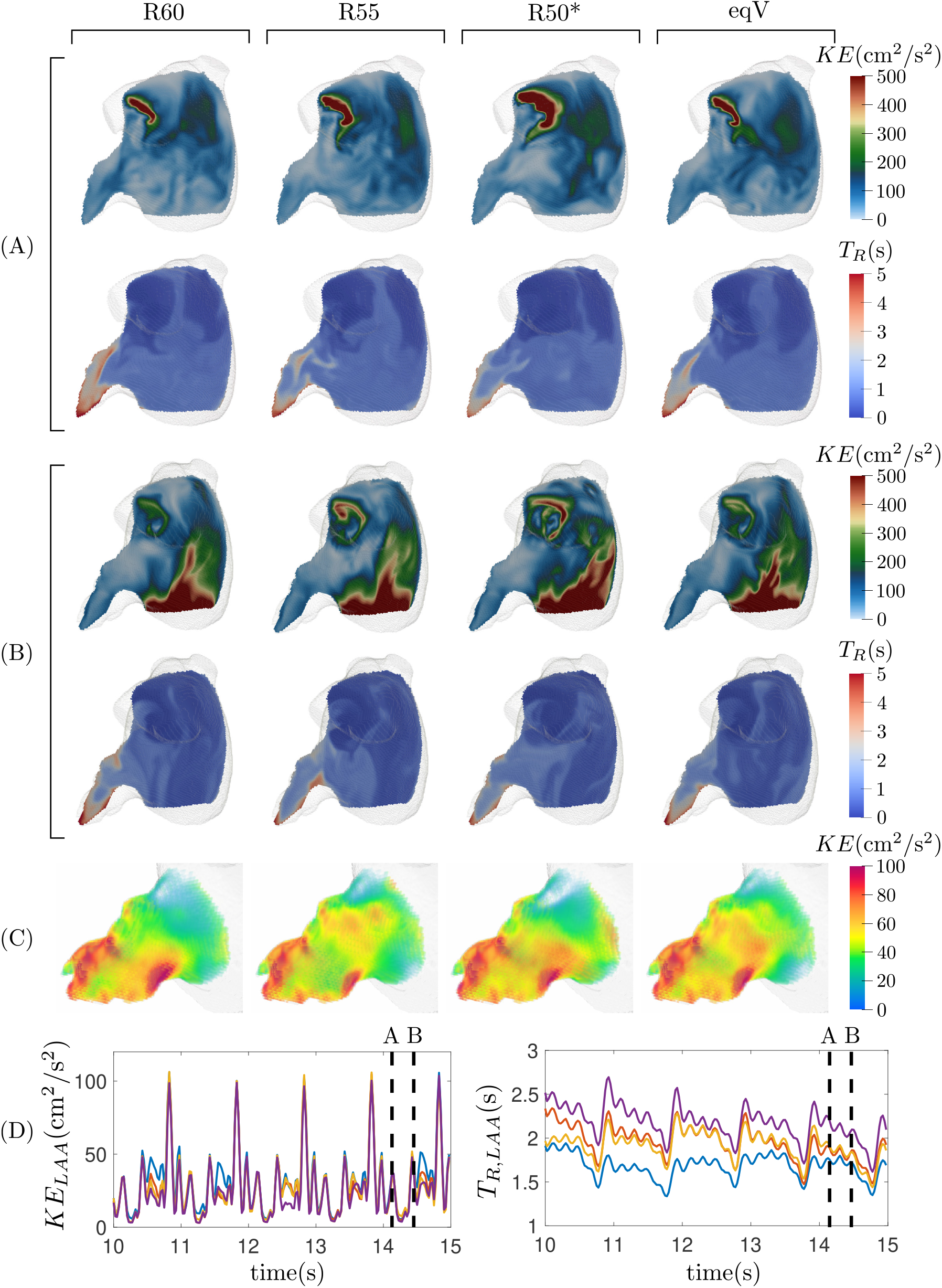
Subject 2. Visualization of the instantaneous kinetic energy and residence time in an oblique plane section of the left atrium for the same two instants (A) and (B) of figure 4. Panel (C) shows the kinetic energy inside the LAA during the ventricle diastole. Panel (D) shows the temporal evolution of the mean kinetic energy and residence time inside the left atrial appendage along five cardiac cycles for the four flow splits (R60, violet; R55, yellow; R50*, blue; eqV, red).

**Fig. 7.**
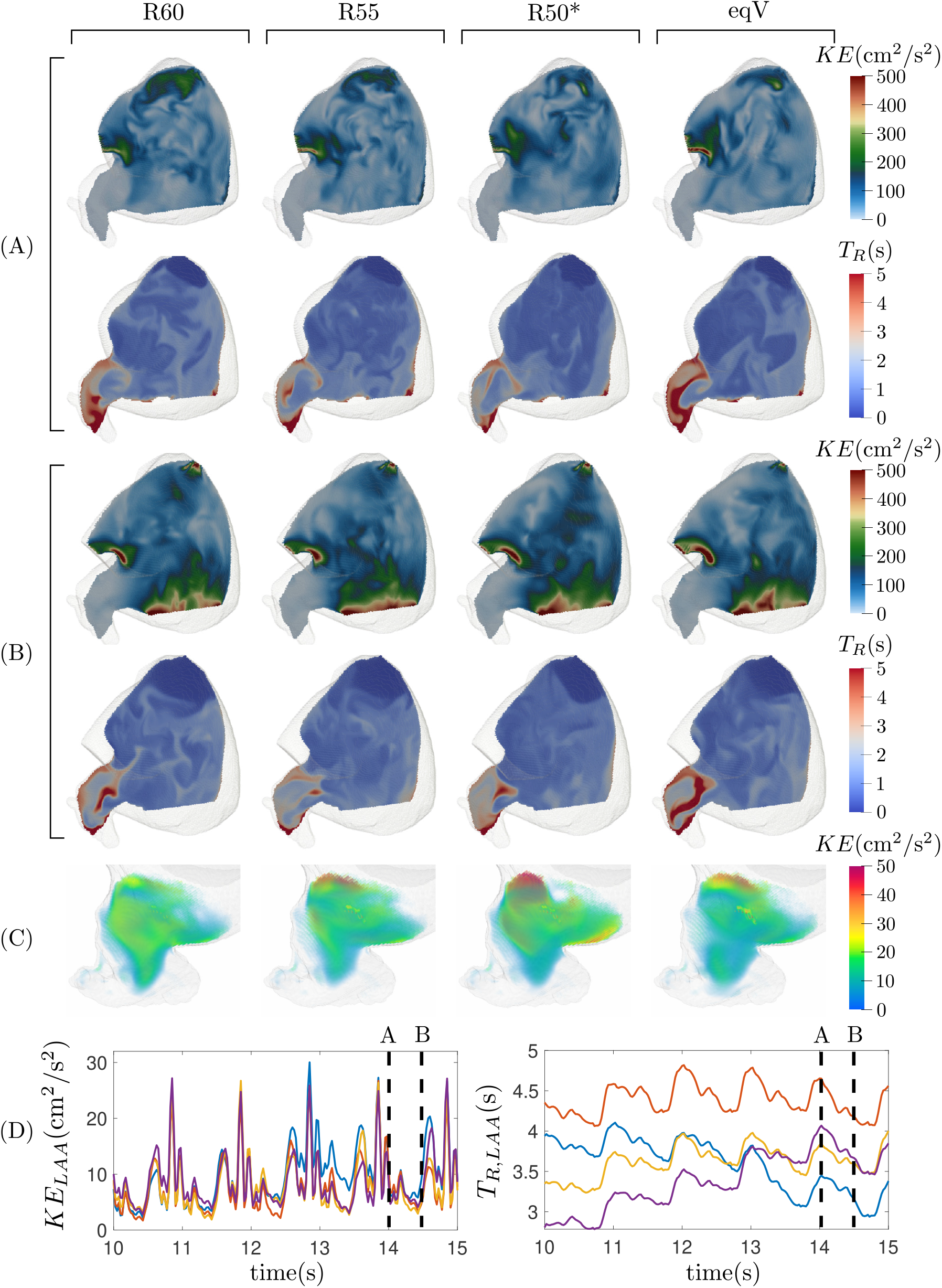
Subject 6. Visualization of the instantaneous kinetic energy and residence time in an oblique plane section of the left atrium for the same two instants (A) and (B) of figure 4. Panel (C) shows the kinetic energy inside the LAA during the ventricle diastole. Panel (D) shows the temporal evolution of the mean kinetic energy and residence time inside the left atrial appendage along five cardiac cycles for the four flow splits (R60, violet; R55, yellow; R50*, blue; eqV, red).

After 13 cardiac cycles, blood residence time inside the left atrium showed stable distributions with small, quasi-periodic variations along the cardiac cycle. The lowest *T_R_* values appear near the PV inlets, where blood enters the chamber. The incoming blood travels toward the mitral valve, washing out the LA body. This clearing process is particularly effective in subjects with normal atrial function, keeping *T_R_* around 1 cycle inside the LA body. But even in subjects with impaired atrial function, the LA body ultimately acts as a conduit, and *T_R_* rarely exceeds 2 cycles in this region of the chamber. Consistent with previous works [24, 37], the *T_R_* is significantly longer inside the LAA than in the LA body. Moreover, and also in contrast to the LA body, the LAA experiences significant variability in these two hemodynamic variables with respect to the PV flow split. For both the normal and impaired, pro-thrombotic atrium, *T_R_* is elevated in the distal LAA regardless of PV flow split; however, the proximal *T_R_* distributions exhibit more variability, especially in the pro-thrombotic case. This variability is captured by computing the spatial average of *T_R_*, which is plotted vs. time together with *KE* in figures 6-7C. These plots confirm that changing the PV flow split alters the *T_R_* inside the LAA all along the cardiac cycle and that, as noted above, these effects are more considerable in the impaired, pro-thrombotic atrium (figure 7) than in the normal one (figure 6). The next sections quantify this variability in more detail across our entire study cohort.

### 3.3 Flow sensitivity to changes in PV flow split

To study in more detail how PV inflow affects blood flow in different regions of the atrium, we analyzed the correlations between flows obtained with different PV flow splits in different LA regions. We first computed the correlation factor of the PV inflow flowrates along the cardiac cycle (*ρ_PV_*, defined in eq.5, summarized in table 2), observing small departures from perfect correlation (*ρ_PV_* = 1) for all cases and low subject-to-subject variability. To investigate the downstream effects of these differences in atrial inflow, we computed correlation factors for the flow velocity inside the atrial body (eq.6) in the directions parallel (*ρ_LAb,||MV_*) and perpendicular (*ρ*_*LAb*,⊥*MV*_) to the MV plane. In this calculation, we excluded the mesh points inside the LAA and the region near the PVs, as indicated in figure 3. These data are summarized in table 2 and plotted in figure 8A-B as functions of *ρ_PV_*. The results suggest that both *ρ_LAb,||MV_* and *ρ*_*LAb*,⊥*MV*_ tend to decrease with *ρ_PV_*, indicating that stronger variations in PV flow split cause stronger variations downstream in the LA body.

**Table 2.**
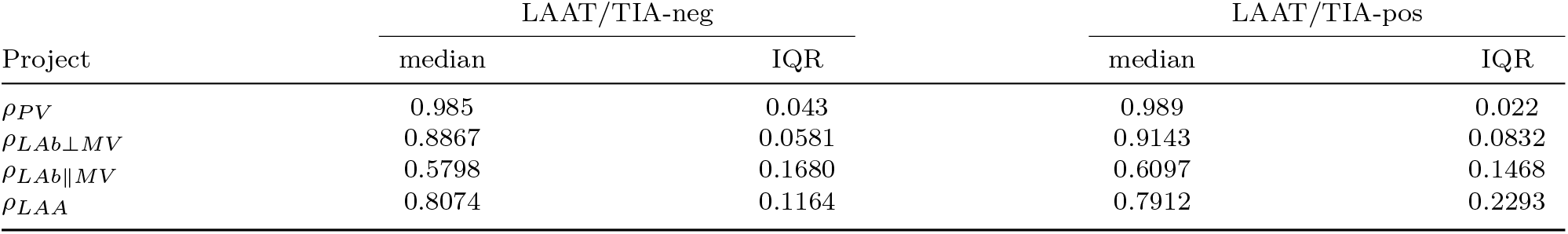
Median and Inter-Quartile Range of the correlation factors of the flow inside the LA body in perpendicular and parallel to the MV plane directions, and the flow inside the LAA for thrombus/TIA negative and positive cases.

**Fig. 8.**
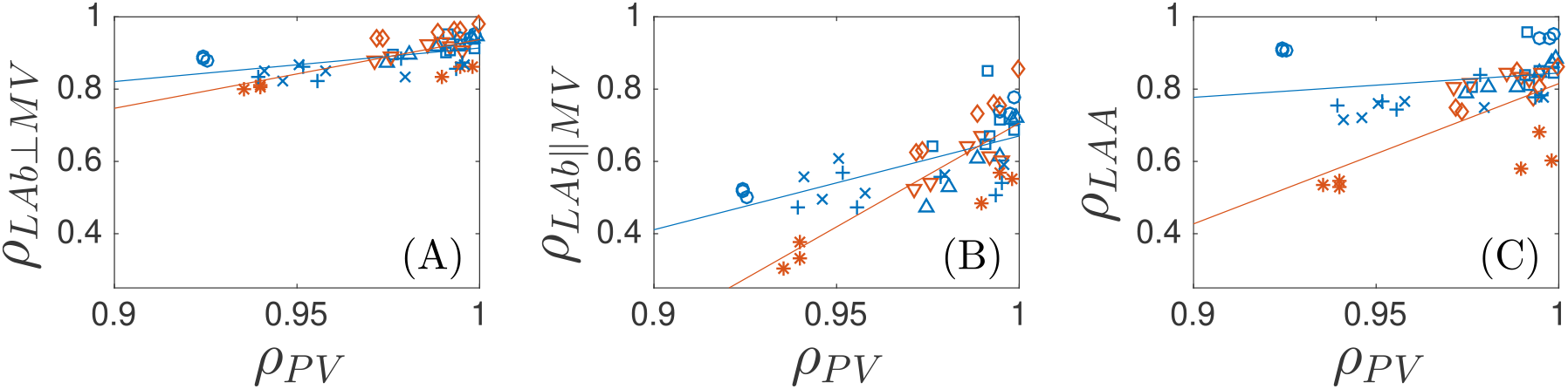
Scatter plot with the correlation factor of the flow through the pulmonary veins versus the correlation factor of the flow inside the left atrium body in perpendicular (A) and parallel (B) directions to the mitral valve and inside the left atrial appendage (C). Symbols represent different cases (from case 1 to 8: ◦, □, △, +, ×, ▽, ◇, *). The straight lines represent the regression lines for thrombus negative (blue) and thrombus positive (red) cases.

We also found that *ρ*_*LAb*,⊥*MV*_ has higher values and a shallower dependence on *ρ_PV_* when compared to *ρ_LAb,||MV_* across all the simulations. These data indicate that variations in PV flowrates have a weaker effect on the main axial transit of blood from the PVs to the MV than on secondary, transverse flow patterns. In the discussion section, we argue that the different sensitivities to PV flow split are related to global chamber mass conservation. Of note, the thrombus/TIA-positive cases exhibit a steeper decrease of *ρ_LAb,||MV_* than the normal cases, suggesting that secondary flow patterns of pro-thrombogenic atria with impaired function (subjects 6–8) are more sensitive to PV inflow flowrates, consistent with the vector velocity, *KE*, and *T_R_* maps presented above. The differences in the sensitivity of the hemodynamics between normal and impaired, prothrombotic atria are particularly noticeable in the LAA, as indicated by the velocity correlation factor *ρ_LAA_* vs. *ρ_PV_* inside the appendage (figure 8C).

### 3.4 Kinetic and residence time statistics in LA body and LAA

Atrial kinetic energy and residence time have been proposed as hemodynamic biomarkers of thrombosis risk [24, 37]. To study how PV flow split variability affects these variables, we plotted their probability box plots using data from three cardiac cycles after converging the simulations (12s < *t* < 15s). figure 9 displays these boxplots for the LA body, showing that the flow in the direction perpendicular to the MV plane (*KE*_⊥_, figure 9A) is more energetic than the secondary flows parallel to that plane (*KE*_||_, figure 9B). Overall, normal atria experienced higher *KE* than impaired, pro-thrombotic ones.

**Fig. 9.**
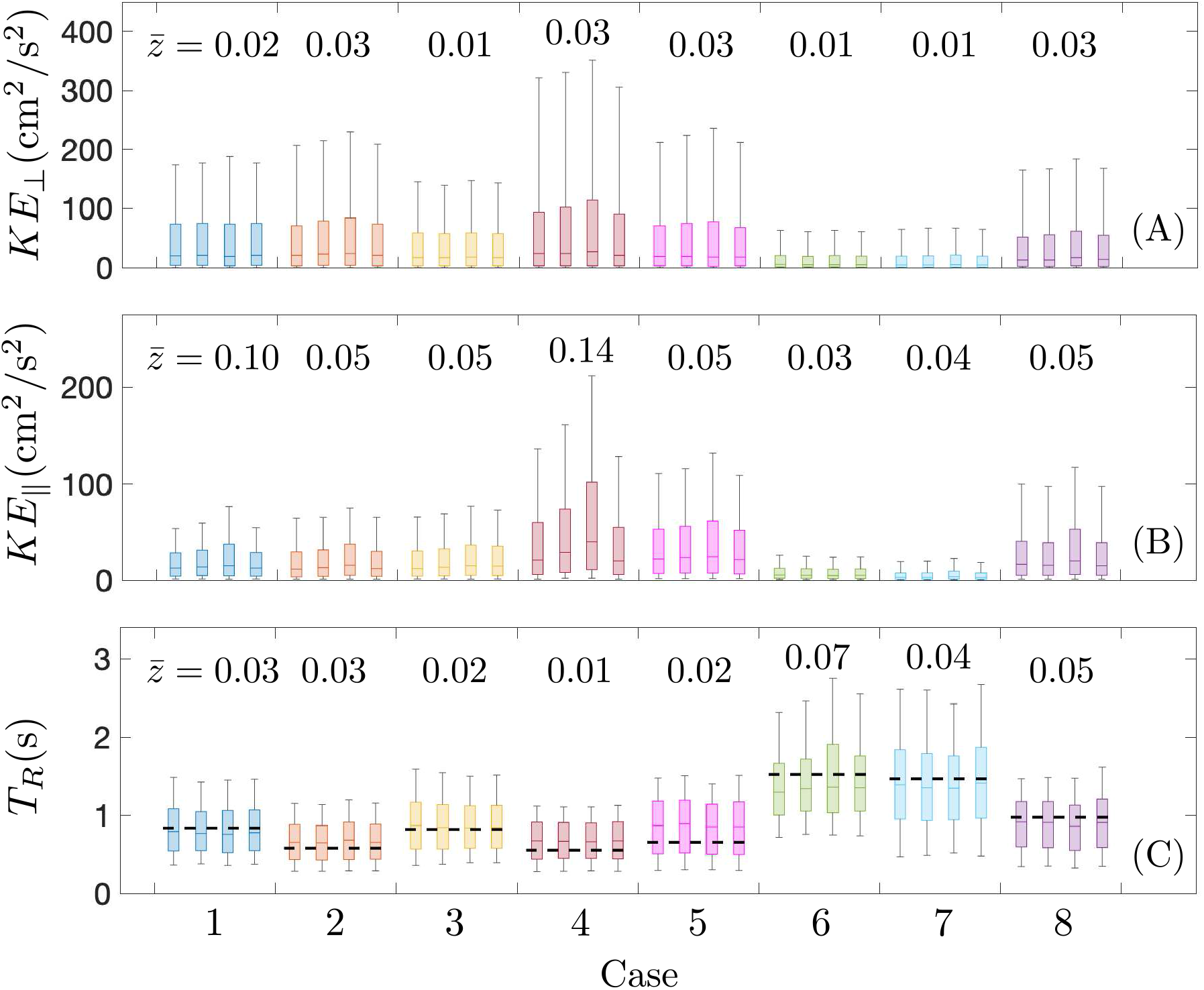
Probability distributions of kinetic energy, in perpendicular (A) and parallel (B) directions to the MV, and residence time (C) inside the left atrium body along three cardiac cycles (12s < *t* < 15s). Boxes and whiskers represent 9, 25, 50, 75 and 91 percentiles for each subject and each flow split condition (from left to right: R60, R55, R50* and eqV). Dashed line represents the residence time prediction given by TR = LAV/(α LVSV), with α = 1.4 and LAV and LVSV are respectively the mean LA volume and LA stroke volume. Mean intra flow split z-scores are represented on the top of each case.

To quantify the level of variability across PV flow splits, we computed z-scores for each patient as *z* = mean 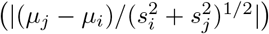, where *μ_i_* is the mean and *s_i_* is the standard deviation of the physical quantity for flow split *i*, and included these data in the figure. Consistent with the correlation factor data reported above, *KE*_⊥_ was less sensitive to the PV flow split than *KE_||_*. Specifically, the z-score medians (range) of *KE*_⊥_ over the normal and pro-thrombotic atria were respectively 0.03 (0.01-0.03) and 0.01 (0.01-0.03), while the corresponding values for *KE*_||_ were 0.05 (0.05-0.14) and 0.04 (0.03-0.05). Interestingly, these z-scores suggest that the flow *KE* of normal and pro-thrombotic atria have similar sensitivities to the PV flow split, which contrast with the results from the correlation factors shown in figure 8. Our interpretation of this seeming discrepancy is that it is the orientation and not the overall magnitude of the velocity field which is more sensitive to PV flow split in the pro-thrombotic atria. It is also worth noting that many of the simulations produce the highest atrial body kinetic energies for the R50* flow split. This peculiarity, which is particularly noticeable for subject 4, can be explained by the differences in PV inlet areas for each subject. The R50* condition splits the PV flowrate evenly among the four PVs, often creating a high inflow velocity through the narrowest PV inlet.

Figure 9C displays barplots of the residence time averaged over 12s < *t* < 15s inside the LA body. Overall, normal atria experienced lower *TR* than impaired, pro-thrombotic ones. Gonzalo et al. [37] suggested that global chamber LA mass conservation and conduit function make *TR* inside the atrial body to be approximated by the simple model *T_R_* = LAV/(*α* LVSV) where LAV and LVSV are respectively the mean LA volume and LV stroke volume, and *α* > 1 is a parameter that accounts for atrial washout occurring mostly during early LV filling. figure 9C confirms the model, which helps understand the low variability of *T_R_* with respect to flow split changes for all the cases. Also, the z-score medians (range) of *T_R_* over the normal and pro-thrombotic atria were respectively 0.02 (0.01-0.03) and 0.05 (0.04-0.07), suggesting that *T_R_* in the body of pro-thrombogenic atria is more sensitive to PV flow split when compared to normal atria.

Figure 10 shows probability box plots of kinetic energy and residence time inside the LAA using data from the same three cardiac cycles (12s < t < 15s), analogous to figure 9. Overall, the magnitude of *KE* in the LAA is significantly lower than the total *KE* in the LA body, although it is comparable to that of atrial body secondary flow patterns (i.e., *KE*_||_). In normal atria, we observe slight differences in LAA KE when the flow split is varied within each subject. These differences become more evident in impaired, pro-thrombotic atria, which, in general, also present smaller LAA *KE* values. Consonantly, the z-scores for LAA *KE* have a median (range) of 0.05 (0.01-0.08) in normal atria vs. 0.1 (0.08-0.14) in pro-thrombotic atria.

**Fig. 10.**
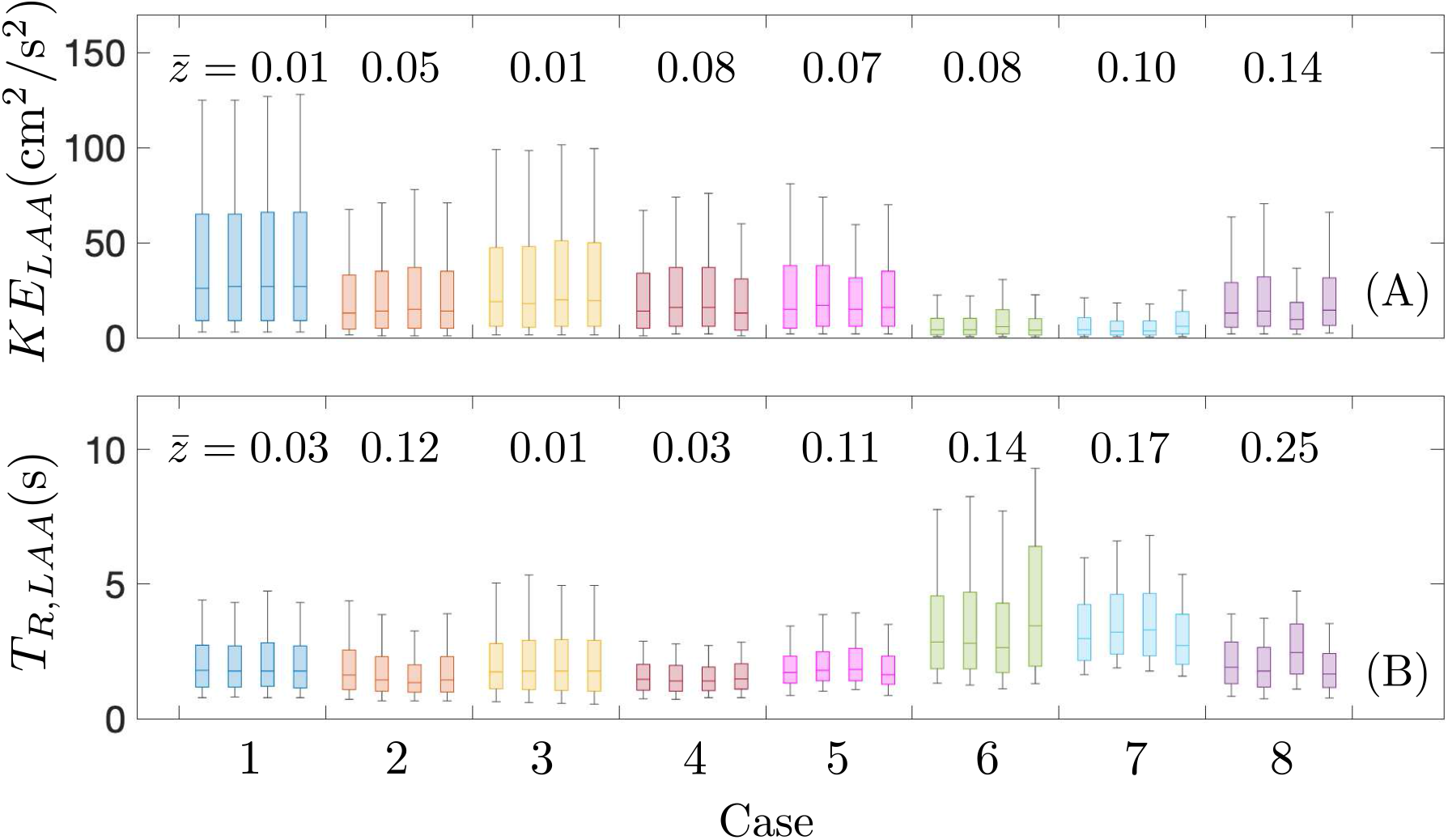
Probability distributions of kinetic energy (A) and residence time (B) inside the left atrial appendage along three cardiac cycles (12s < *t* < 15s). Boxes and whiskers represent 9, 25, 50, 75 and 91 percentiles for each subject and flow split condition (from left to right: R60, R55, R50* and eqV). Mean intra flow split z-scores are represented on the top of each case.

Residence time inside the LAA is significantly higher than inside the LA body, specially its maximum values, which often correspond with stagnant blood pools in the LAA apex (see, e.g., figures 6 and 7 and refs [24, 37]). In normal atrial, the median (range) of the LAA *T_R_ z*-scores is 0.03 (0.01 - 0.12), and two of the five cases have z-scores > 0.1. In pro-thrombotic atria, the variability induced by PV flow split in LAA *T_R_* was more pronounced, with z-score statistics being 0.17 (0.13 - 0.25).

The variability in LAA residence time induced by the PV flow split could be a significant source of uncertainty when using *T_R_* to identify patients at risk of atrial thrombosis. To evaluate this possibility, we plotted the mean LAA residence time (12s < *t* < 15s) vs. PV flow split condition for this study’s normal (figure 11A) and impaired, pro-thrombotic (figure 11 B) atria. These plots reveal that the normal atria with highest *T_R_* values overlap with the envelope of the pro-thrombotic cases (red shaded patch in figure 11A), while the pro-thrombotic atrium with lowest *T_R_* overlaps with the envelope of the normal cases (blue shaded patch in figure 11B). These overlaps are primarily caused by the significant sensitivity to PV flow split of the residence time in the pro-thrombotic LAA.

**Fig. 11.**
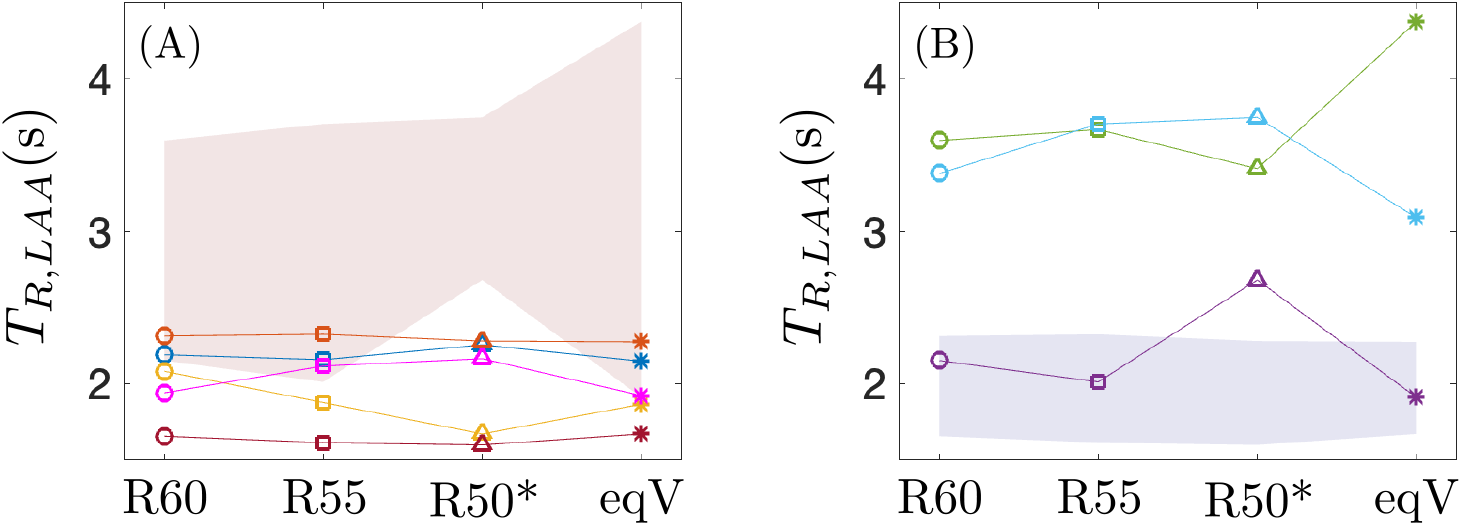
Mean value of the residence time in the left atrial appendage along three cardiac cycles (12s < *t* < 15s) for the four flow splits. Data in (A) correspond to the thrombus-negative cases, and the shaded patch in red corresponds to the envelope of the thrombus-positive cases. Data in (B) correspond to the thrombus-positive cases, and the shaded patch in blue corresponds to the thrombus-negative cases. The colors of the subjects are the same as in figures 8 and 9.

## 4 Discussion

The past decade has witnessed significant growth in the number of computational fluid dynamics (CFD) analyses of the left atrium [17, 18, 24, 25, 30, 38, 53]. These works shed light onto the anatomical and physiological determinants of atrial flow and offered proof-of-principle validation for blood stasis mapping as a thrombosis predictor. In parallel, echocardiographic LV stasis mapping has proven predictive value for LV thrombosis and brain embolism in pre-clinical and clinical pilot studies [54–56]. However, blood flow in the LA is highly three-dimensional and unsteady, and CFD-simulated LA hemodynamics can exhibit significant intra-patient variability depending on modeling parameters [17, 21, 36, 53]. A better understanding of this variability is required to develop reliable CFD-derived biomarkers of atriogenic stroke risk.

### 4.1 Pulmonary vein inflow, atrial hemodynamics, and LAA stasis

The LA contributes to cardiac pumping via three distinct functions. When the mitral valve is closed, it receives flow kinetic energy from the lungs and transforms it into myocardial elastic energy that is released back into the flow during early LV filling (reservoir function). It actively contracts to drive late LV filling (booster function). And, while the mitral valve is open, it forms a tube through which the pulmonary veins continuously discharge into the LV (conduit function) [57]. These three functions involve flow interactions between the atrium and the pulmonary veins and can overlap during some phases of the cardiac cycle, creating complex PV inflow profiles with several forward waves during atrial and early LV diastole, and alternating forward and backward waves during atrial systole. There are usually two pairs of PV inlets in the LA, the right pair opposes the left pair creating two opposing inflow jets. Furthermore, the precise number and anatomical configuration of PV inlets can vary significantly among patients [25].

These factors give rise to intricate atrial flow patterns, first visualized in 3D approximately two decades ago [58, 59]. However, it has not been until recently that the influence of PV inflow in LA hemodynamics began to be investigated in detail using CFD simulations [25, 29, 42]. Lantz et al.[42] performed 4D flow CMR measurements and CT-based CFD simulations of the whole left heart, varying the PV flow split. A strength of this study was its design of experiments (DOE) approach to interrogate the three-dimensional parameter space defined by varying the PV flow split of the four pulmonary veins. Following this systematic approach, Lantz et al.[42] ran 20 simulations per patient in N=3 patients, which provided unprecedented information on the PV inflow dependence of LA and LV flow. They reported a large variability for the LA body *KE*, although this may be caused by the fact that some of the PV flow splits considered in the DOE analysis were unphysiological. On the other hand, their 25-25-25-25 simulations, which are equivalent to our R50* condition, yielded LA *KE* profiles in the middle of the DOE envelope and were in fairly good agreement with the 4D flow MRI patient-specific measurements. Unfortunately, Lantz et al.[42] did not compute residence time or report flow data inside the LAA, the most frequent site of intracardiac thrombosis [60].

More recently, Mill et al.[25] performed an exhaustive investigation of the influence of atrial anatomical parameters on LA hemodynamics and LAA stasis. Their approach was complementary to that of Lantz et al.[42]; they considered a large cohort of atrial fibrillation patients (N=52, 25 of them thrombotic), performing one simulation per patient using the inflow/outflow profiles for all simulations and computing a comprehensive set of hemodynamic descriptors in the LAA. Then, they interrogated the database for correlations between anatomical features and hemodynamic descriptors across different patients. Of particular interest for the present study, Mill et al.[25] analyzed the orientation of the PV inlets, finding that there is significantly more variability in the right-side PVs than in the left-side PVs, and that this variability significantly affected atrial hemodynamics. Nevertheless, Mill et al.[25] did not find a clear association between right PV orientations and LAA thrombosis. Likewise, they found no direct link between LAA thrombosis and the number of PVs, emphasizing the multi-factorial nature of LAA stasis

Thus, the existing literature suggests that the effects of the PV inflow profiles on atrial hemodynamics are significant and may be difficult to disentangle from anatomical factors. Furthermore, despite these recent works, there is a paucity of data on how the PV flow split influences LAA stasis, a major determinant of atrial thrombosis associated with stroke risk. This paucity creates a significant problem because the PV flow split is a crucial parameter to define inflow boundary conditions and its patient-specific measurement is challenging [39]. To shed light on this question, we performed a CFD investigation on a moderate-size cohort (N=8) while also considering four different PV flow split scenarios per patient-specific anatomical model. While our study did not perform an exhaustive DOE parametric analysis [42] or considered a large cohort [25], it balanced intersubject and intra-subject variability by including normal and thrombotic atria, and by focusing on relatively small, physiologically relevant departures from the 25-25-25-25 flow split. In addition, we conducted high-fidelity CFD simulations in terms of spatial and temporal resolution (see, e.g., ref [61]), the number of cycles run, and computed high-cost hemodynamic variables like blood residence time, which are tightly linked with blood stasis [62–65], and have been clinically validated as thrombosis biomarkers in other cardiac chambers [54–56].

In agreement with the previous studies, instantaneous visualizations of our CFD simulations show that the PV flow split modifies the flow patterns in the LA and LAA. While this effect is complex and multifactorial, our analysis suggests a few general trends. First, the velocity along the PV-MV axis delineating the atrial conduit route (denoted as the ⊥component) is relatively insensitive to the PV flow split. A possible explanation for this behavior is that the flow rate through the MV is dictated by global mass conservation of the LA chamber independent of the flow split, and this flow rate, in turn, dominates fluid motion along the PV-MV axis. Consistent with this hypothesis, Lantz et al.[42] reported CFD-derived flow visualizations in the MV plane that were relatively insensitive to PV flow split and agreed well with 4D flow MRI measurements. Also consistent with this hypothesis, our simulations show that fluid motions parallel to the MV plane (denoted as the || component), which are not constrained by global LA mass conservation, are more sensitive to the PV flow split. Since the flow velocities along the PV-MV axis are stronger, this effect is most apparent when separately computing the flow kinetic energy in the || and ⊥ components. Blood residence time in the atrial body, which is primarily dictated by the ⊥ component, is relatively insensitive to the PV flow split. The sensitivity of LAA hemodynamics to the PV flow split is more involved and is discussed below.

### 4.2 The hemodynamic sensitivity of the left atrial appendage

The LAA is a small tubular sac protruding from the atrial body in a direction roughly perpendicular to the main transit path of LA flow between the PVs and the MV. Consequently, blood residence time inside the LAA significantly exceeds that in the atrial body [17]. Given its anatomical isolation from the conduit that joins the PVs and the MV, one could intuit that LAA hemodynamics should be relatively insensitive to the PV flow split. However, our results indicate that blood flow and, in particular blood residence time, are more sensitive to the PV flow split in the LAA than in the atrial body. As we argued above, secondary flow patterns perpendicular to the PV-MV axis (i.e., the || component) are not significantly constrained by global chamber mass conservation and, thus, could vary appreciably with anatomical parameters, boundary conditions (including the PV flow split), blood rheology (e.g., non-Newtonian effects [37]), etc. Our data suggest that changes in secondary flow patterns near the LAA ostium can affect flow inside the LAA and, as a result, the LAA residence time. This idea is supported by Garcia-Isla et al’s data showing that perturbations to the PV diameter that increase blood velocity at the ostium also decrease the endothelial cell activation potential [29]. This mechanism is highly dependent on each patient’s LAA anatomical features and its specific orientation with respect to the PV inlets and the PV-MV axis [25], which would explain why some of our subjects exhibit small variations in mean LAA *T_R_* with PV flow split (i.e., cases 1-3), while others exhibit large enough variations to create overlap between the thrombus/TIA group and the normal group. Unlike the atrial body, the appendage is a closed chamber with no conduit function that relies primarily on the expansion and contraction of its walls for blood clearance. This mechanism is quantified by the LAA emptying fraction, (LAA EF, Table 1), which reached 0.53 +/- 0.08 (mean +/- std) in this study’s normal atria and 0.21 +/- 0.01 in the impaired, prothrombotic atria. The significant difference in LAA EF between groups is congruent with the higher TR found in the impaired prothrombotic atria and, relevant to the present study, could also explain the higher variability of this group’s LAA *T_R_* with the PV flow split. This may be because secondary swirling motions with zero net volume flux play a more important role in washing out LAAs of lower emptying fraction, and these fluid motions are more sensitive to anatomical factors and PV flow profiles [37]. Consistent with this hypothesis, we recently showed that the LAA *T_R_* of normal atria is less sensitive to non-Newtonian effects than the LAA TR of impaired prothrombotic atria [37]. Finally, we note that we cannot discard the alternate hypothesis that LA EF and not LAA EF drives the variability of hemodynamic variables, since the two EFs are highly correlated in our cohort (correlation coefficient = 0.90, p=0.002).

### 4.3 Study Limitations

The cohort used in this study is small (N=8). This is partly due to the computational cost of running high-resolution CFD simulations for 15 cycles at *CFL* < 0.3 and the need to run four simulations per subject to study the effect of the PV flow split. However, high temporal resolution is crucial to obtain convergence in hemodynamic metrics obtained from CFD analysis of LA hemodynamics [61]. To help compensate for the low N, we selected a diverse cohort in terms of LA volume and function (see Table 1). While this strategy allowed us to cover a wide range of patient-specific conditions, it might bias our observations regarding inter-patient vs. intra-patient variability. In addition, all of the 8 patients in this study had normal pulmonary vein anatomy with 2 right and 2 left pulmonary veins. Whether the results extrapolate to anatomic variants was not evaluated.

We consider PV flow splits in the ranges (left/right) 40-50%/50-60% to compare differences in LA hemodynamics between the even split [7, 17, 24, 37, 66, 67] and equal velocity inflow conditions [19, 21, 34, 68, 69] often used in simulation studies, and the more physiological uneven splits caused by the right lung being larger than the left lung. However, multiple factors can alter the PV flow split outside of the range we studied, including common ones like changes in body position [41] and less common ones like abnormalities in the pulmonary artery or veins [70, 71], or lung resection surgeries [72]. Additionally, body position can also affect superior/inferior PVs flow split, whose effect would be interesting to study in future work.

All our simulations were run at a constant heart rate of 60 min^-1^ corresponding to resting conditions and constant (i.e., Newtonian) viscosity. This choice is customary [21, 24, 30, 61, 67, 73] and justified based on the rationale of reducing the number of independent parameters to the PV flow split. The flow split has been reported to change with heart rate, approaching 50-50% during exercise [74], and heart rate could alter blood viscosity inside the LA via non-Newtonian effects [37]. In turn, non-Newtonian effects could be interlinked with flow-split-induced variability in the LAA, where long residence times contribute to increased blood viscosity [37].

## 5 Conclusion

The pulmonary vein flow split is an important parameter governing boundary conditions in patient-specific left atrial blood flow simulations. This work investigated how the PV flow split influences left atrial flow and LAA stasis by performing CFD simulations on subjects with normal and impaired pro-thrombotic atria. We found that secondary flow patterns in planes parallel to the mitral valve are sensitive to variations in PV flow split. Flow inside the LAA, particularly blood residence time, is also affected by the changes in PV flow split. This sensitivity is highest in patients with reduced LAA emptying fraction, highlighting the need for accounting for uncertainty in CFD-derived risk scores of atrial thrombosis.

## Acknowledgments

This work was partially supported by Comunidad de Madrid (Synergy Grant Y2018/BIO-4858 PREFI-CM), Spanish Research Agency(AEI, grant number PID2019-107279RB-I00), Institute de Salud Carlos III (grant numbers PI15/02211-ISBITAMI and DTS/1900063-ISBIFLOW), and by the EU—European Regional Development Fund.

## References

[1] Benjamin, E.J., Muntner, P., Alonso, A., et al.: Heart disease and stroke statistics-2019 update: A report from the American Heart Association. Circulation 139(10), 56–528 (2019) https://doi.org/10.1161/CIR.0000000000000659

[2] Kamel, H., P.M., O., Elkind, M.S.V., Iadecola, C.: Atrial fibrillation and mechanisms of stroke: time for a new model. Stroke 47(3), 895–900 (2016) https://doi.org/10.1161/STROKEAHA.115.012004

[3] Cresti, A., García-Fernandez, M.A., Sievert, H., Mazzone, P., et al.: Prevalence of extra-appendage thrombosis in non-valvular atrial fibrillation and atrial flutter in patients undergoing cardioversion: a large transoesophageal echo study. EuroIntervention 15(3), 225–230 (2019) https://doi.org/10.4244/EIJ-D-19-00128

[4] Aakre, C.A., McLeod, C.J., Cha, S.S., Tsang, T.S.M., et al.: Comparison of clinical risk stratification for predicting stroke and thromboembolism in atrial fibrillation. Stroke 45(2), 426–431 (2014) https://doi.org/10.1161/STROKEAHA.113.002585

[5] Siddiqi, T.J., Usman, M.S., Shahid, I., Ahmed, J., et al.: Utility of the CHA2DS2-VASc score for predicting ischaemic stroke in patients with or without atrial fibrillation: a systematic review and meta-analysis. Eur. J. Prev. Cardiol. 29(4), 625–631 (2022) https://doi.org/10.1093/eurjpc/zwab018

[6] Mihalef, V., Ionasec, R.I., Sharma, P., Georgescu, B., et al.: Patient-specific modelling of whole heart anatomy, dynamics and haemodynamics from four-dimensional cardiac CT images. Interface Focus 1(3), 286–296 (2011) https://doi.org/10.1098/rsfs.2010.0036

[7] Chnafa, C., Mendez, S., Nicoud, F.: Image-based large-eddy simulation in a realistic left heart. Comput. Fluids 94, 173–187 (2014) https://doi.org/10.1016/j.compfluid.2014.01.030

[8] Vedula, V., George, R., Younes, L., Mittal, R.: Hemodynamics in the left atrium and its effect on ventricular flow patterns. J. Biomech. Eng. 137(11), 111003 (2015) https://doi.org/10.1115/1.4031487

[9] Lantz, J., Henriksson, L., Persson, A., Karlsson, M., Ebbers, T.: Patient-specific simulation of cardiac blood flow from high-resolution computed tomography. J. Biomech. Eng. 138(12) (2016) https://doi.org/10.1115/1.4034652

[10] Bucelli, M., Zingaro, A., Africa, P.C., Fumagalli, I., et al.: A mathematical model that integrates cardiac electrophysiology, mechanics, and fluid dynamics: Application to the human left heart. Int J Numer Meth Biomed Engng 39(3) (2023) https://doi.org/10.1002/cnm.3678

[11] Watanabe, H., Sugiura, S., Kafuku, H., Hisada, T.: Multiphysics simulation of left ventricular filling dynamics using fluid-structure interaction finite element method. Biophys. J. 87(3), 2074–2085 (2004) https://doi.org/10.1529/biophysj.103.035840

[12] Seo, J.H., Vedula, V., Abraham, T., Lardo, A.C., et al.: Effect of the mitral valve on diastolic flow patterns. Phys. Fluids 26(12) (2014) https://doi.org/10.1063/1.4904094

[13] Arefin, M.S., Morsi, Y.S.: Fluid structure interaction (FSI) simulation of the left ventricle (LV) during the early filling wave (E-wave), diastasis and atrial contraction wave (A-wave). Australas. Phys. Eng. Sci. Med. 37(2), 413–423 (2014) https://doi.org/10.1007/s13246-014-0250-4

[14] Vedula, V., Seo, J., Lardo, A.C., Mittal, R.: Effect of trabeculae and papillary muscles on the hemodynamics of the left ventricle. Theor. Comput. Fluid Dyn. 30, 3–21 (2016) https://doi.org/10.1007/s00162-015-0349-6

[15] Prisco, A.R., Aliseda, A., Beckman, J.A., Mokadam, N.A., et al.: Impact of LVAD implantation site on ventricular blood stagnation. ASAIO J. 63(4), 392–400 (2018) https://doi.org/10.1097/MAT.0000000000000503

[16] Liao, S., Neidlin, M., Li, Z., Simpson, B., Gregory, S.D.: Ventricular flow dynamics with varying LVAD inflow cannula lengths: In-silico evaluation in a multiscale model. J. Biomech. 72, 106–115 (2018) https://doi.org/10.1016/j.jbiomech.2018.02.038

[17] Koizumi, R., Funamoto, K., Hayase, T., Kanke, Y., et al.: Numerical analysis of hemodynamic changes in the left atrium due to atrial fibrillation. J. Biomech. 48(3), 472–478 (2015) https://doi.org/10.1016/j.jbiomech.2014.12.025

[18] Otani, T., Al-Issa, A., Pourmorteza, A., McVeigh, E.R., et al.: A computational framework for personalized blood flow analysis in the human left atrium. Ann. Biomed. Eng. 44(11), 3284–3294 (2016) https://doi.org/10.1007/s10439-016-1590-x

[19] Masci, A., Alessandrini, M., Forti, D., Menghini, F., et al.: A proof of concept for computational fluid dynamic analysis of the left atrium in atrial fibrillation on a patient-specific basis. J. Biomech.Eng. 142(1), 011002 (2020) https://doi.org/10.1115/1.4044583

[20] Dueñas-Pamplona, J., García García, J., Sierra-Pallares, J., Ferrera, C., et al.: A comprehensive comparison of various patient-specific CFD models of the left atrium for atrial fibrillation patients. Comput. Biol. Med. 133, 104423 (2021) https://doi.org/10.1016/j.compbiomed.2021.104423

[21] Corti, M., Zingaro, A., Dede, L., Quarteroni, A.: Impact of atrial fibrillation on left atrium haemodynamics: A computational fluid dynamics study. Comput. Biol. Med. 150, 106143 (2022) https://doi.org/10.1016/j.compbiomed.2022.106143

[22] Al-Saady, N.M., Obel, O.A., Camm, A.J.: Left atrial appendage: structure, function, and role in thromboembolism. Heart 82(5), 547–554 (1999) https://doi.org/10.1136/hrt.82.5.547

[23] Leventić, H., Babin, D., Velicki, L., Devos, D., et al.: Left atrial appendage segmentation from 3D CCTA images for occluder placement procedure. Comput. Biol. Med. 104, 163–174 (2019) https://doi.org/10.1016/j.compbiomed.2018.11.006

[24] García-Villalba, M., Rossini, L., Gonzalo, A., Vigneault, D., et al.: Demonstration of patient-specific simulations to assess left atrial appendage thrombogenesis risk. Front. Physiol. 12, 596596 (2021) https://doi.org/10.3389/fphys.2021.596596

[25] Mill, J., Harrison, J., Legghe, B., Olivares, A.L., et al.: In-silico analysis of the influence of pulmonary vein configuration on left atrial haemodynamics and thrombus formation in a large cohort. In: Functional Imaging and Modeling of the Heart, pp. 605–616 (2021). https://doi.org/10.1007/978-3-030-78710-3_58

[26] Di Biase, L., Santangeli, P., Anselmino, M., Mohanty, P., et al.: Does the left atrial appendage morphology correlate with the risk of stroke in patients with atrial fibrillation? Results from a multicenter study. J. Am. Coll. Cardiol. 60(6), 531–538 (2012) https://doi.org/10.1016/j.jacc.2012.04.032

[27] Bisbal, F., Guiu, E., Calvo, N., Marin, D., et al.: Left atrial sphericity: A new method to assess atrial remodeling. Impact on the outcome of atrial fibrillation ablation. J. Cardiovasc. Electrophysiol. 24(7), 752–759 (2013) https://doi.org/10.1111/jce.12116

[28] Yamamoto, M., Seo, Y., Kawamatsu, N., Sato, K., et al.: Complex left atrial appendage morphology and left atrial appendage thrombus formation in patients with atrial fibrillation. Circ. Cardiovasc. Imaging 7(2), 337–343 (2014) https://doi.org/10.1161/CIRCIMAGING.113.001317

[29] Garcia-Isla, G., Olivares, A.L., Silva, E., Nuñez-Garcia, M., et al.: Sensitivity analysis of geometrical parameters to study haemodynamics and thrombus formation in the left atrial appendage. Int. J. Numer. Method. Biomed. Eng., 3100 (2018) https://doi.org/10.1002/cnm.3100

[30] Masci, A., Barone, L., Dede, L., Fedele, M., et al.: The impact of left atrium appendage morphology on stroke risk assessment in atrial fibrillation: A computational fluid dynamics study. Front. Physiol 9, 1938 (2019) https://doi.org/10.3389/fphys.2018.01938

[31] Yaghi, S., Chang, A.D., Akiki, R., Collins, S., et al.: The left atrial appendage morphology is associated with embolic stroke subtypes using a simple classification system: A proof of concept study. J. Cardiovasc. Comput. Tomogr. 14(1), 27–33 (2020) https://doi.org/10.1016/j.jcct.2019.04.005

[32] Dueñas-Pamplona, J., García García, J., Castro, F., Muñoz-Paniagua, J., et al.: Morphing the left atrium geometry: A deeper insight into blood stasis within the left atrial appendage. Appl. Math. Model. 108(10), 27–45 (2022) https://doi.org/10.1016/j.apm.2022.03.012

[33] Feng, L., Gao, H., Griffith, B., Niederer, S., Luo, X.: Analysis of a coupled fluid-structure interaction model of the left atrium and mitral valve. Int. J. Numer. Method. Biomed. Eng. 35(11), 3254 (2019) https://doi.org/10.1002/cnm.3254

[34] Qureshi, A., Darwish, O., Dillon-Murphy, D., Chubb, H., et al.: Modelling left atrial flow and blood coagulation for risk of thrombus formation in atrial fibrillation. In: 2020 Computing in Cardiology, pp. 1–4 (2020). https://doi.org/10.22489/CinC.2020.219

[35] Meschini, V., Viola, F., Verzicco, R.: Heart rate effects on the ventricular hemodynamics and mitral valve kinematics. Comput. Fluids 197, 104359 (2020) https://doi.org/10.1016/j.compfluid.2019.104359

[36] Gonzalo, A., Augustin, C., García-Villalba, M., Martinez-Legazpi, P., et al.: Multi-physics, patient-specific computational modeling of LA electrophysiology, biomechanics and hemodynamics reveals mechanistic connections between fibrotic remodeling and thrombosis risk. Heart Rhythm 18(8), 372–373 (2021) https://doi.org/10.1016/j.hrthm.2021.06.925

[37] Gonzalo, A., García-Villalba, M., Rossini, L., Durán, E., et al.: Non-Newtonian blood rheology impacts left atrial stasis in patient-specific simulations. Int. J. Numer. Method Biomed. Eng. 38(6), 3597 (2022) https://doi.org/10.1002/cnm.3597

[38] Paliwal, N., Ali, R.L., Salvador, M., O’Hara, R., et al.: Presence of left atrial fibrosis may contribute to aberrant hemodynamics and increased risk of stroke in atrial fibrillation patients. Front. Physiol. 12, 657452 (2021) https://doi.org/10.3389/fphys.2021.657452

[39] Fadel, B.M., Pibarot, P., Kazzi, B.E., Mahjoub, H., et al.: Spectral Doppler interrogation of the pulmonary veins for the diagnosis of cardiac disorders: A comprehensive review. J. Am. Soc. Echocardiogr. 34(3), 223–236 (2020) https://doi.org/10.1016/j.echo.2020.09.012

[40] Cheng, C.P., Taur, A.S., Lee, G.S., Goris, M.L., Feinstein, J.A.: Relative lung perfusion distribution in normal lung scans: Observations and clinical implications. Congenit Heart Dis. 1(5), 210–216 (2006) https://doi.org/10.1111/j.1747-0803.2006.00037.x

[41] Wieslander, B., Ramos, J.G., Ax, M., Petersson, J., Ugander, M.: Supine, prone, right and left gravitational effects on human pulmonary circulation. J. Cardiovasc. Magn. Reson. 21(1), 69 (2019) https://doi.org/10.1186/s12968-019-0577-9

[42] Lantz, J., Gupta, V., Henriksson, L., Karlsson, M., et al.: Impact of pulmonary venous inflow on cardiac flow simulations: Comparison with in vivo 4D flow MRI. Ann. Biomed. Eng. 74(2), 413–424 (2019) https://doi.org/10.1007/s10439-018-02153-5

[43] Otani, T., Yoshida, T., Yi, W., Endo, S., Wada, S.: On the impact of left upper lobectomy on the left atrial hemodynamics. Front. Physiol. 13, 830436 (2022) https://doi.org/10.3389/fphys.2022.830436

[44] Yi, W., Otani, T., Yoshida, T., Endo, S., Wada, S.: Computational study on hemodynamic effects of left superior pulmonary vein resection and associated physiological changes in the left atrium after left upper lobectomy. Comput. Methods Biomech. Biomed. Engin., 1–12 (2023) https://doi.org/10.1080/10255842.2023.2178258

[45] Yushkevich, P.A., Piven, J., Hazlett, H.C., Smith, R.G., et al.: User-guided 3D active contour segmentation of anatomical structures: significantly improved efficiency and reliability. Neuroimage 31(3), 1116–1128 (2006) https://doi.org/10.1016/j.neuroimage.2006.01.015

[46] Qianqian, F., Boas, D.A.: Tetrahedral mesh generation from volumetric binary and grayscale images. In: 2009 IEEE International Symposium on Biomedical Imaging: From Nano to Macro, pp. 1142–1145 (2009). https://doi.org/10.1109/ISBI.2009.5193259

[47] Myronenko, A., Song, X.: Point set registration: coherent point drift. IEEE Trans. Pattern Anal. Mach. Intell. 32(12), 2262–2275 (2010) https://doi.org/10.1109/TPAMI.2010.46

[48] Otani, T., Shiga, M., Endo, S., Wada, S.: Performance assessment of displacement-field estimation of the human left atrium from 4D-CT images using the coherent point drift algorithm. Comput. Biol. Med. 114, 103454 (2019) https://doi.org/10.1016/j.compbiomed.2019.103454

[49] Hirsch, C.: Numerical Computation of Internal and External Flows. Butterworth-Heinemann, United Kingdom (2007)

[50] Ho, S.Y., Cabrera, J.A., Sanchez-Quintana, D.: Left atrial anatomy revisited. Circ. Arrhythm. Electrophysiol. 5(1), 220–228 (2012) https://doi.org/10.1161/CIRCEP.111.962720

[51] Rossini, L., Martinez-Legazpi, P., Vu, V., Fernandez-Friera, L., et al.: A clinical method for mapping and quantifying blood stasis in the left ventricle. J. Biomech. 49(11), 2152–2161 (2016) https://doi.org/10.1016/j.jbiomech.2015.11.049

[52] Shu, C.W.: Essentially non-oscillatory and weighted essentially non-oscillatory schemes for hyperbolic conservation laws. In: Quarteroni, A. (ed.) Advanced Numerical Approximation of Nonlinear Hyperbolic Equations. Lecture Notes in Mathematics, pp. 325–432. Springer, Berlin (1992)

[53] Bosi, G.M., Cook, A., Rai, R., Menezes, L.J., et al.: Computational fluid dynamic analysis of the left atrial appendage to predict thrombosis risk. Front. Cardiovasc. Med. 5(34) (2018) https://doi.org/10.3389/fcvm.2018.00034

[54] Martinez-Legazpi, P., Rossini, L., Villar, C., Benito, Y., et al.: Stasis mapping using ultrasound: A prospective study in acute myocardial infarction. JACC Cardiovasc. Imaging. 11(3), 514–515 (2018) https://doi.org/10.1016/j.jcmg.2017.06.012

[55] Delgado-Montero, A., Martinez-Legazpi, P., Desco, M.M., Rodríguez-Pérez, D., et al.: Blood stasis imaging predicts cerebral microembolism during acute myocardial infarction. J. Am. Soc. Echocardiogr. 33(3), 389–398 (2020) https://doi.org/10.1016/j.echo.2019.09.020

[56] Rodríguez-González, E., Martínez-Legazpi, P., Mombiela, T., González-Mansilla, A., et al.: Stasis imaging predicts the risk of cardioembolic stroke related to acute myocardial infarction. Eur. Heart J. Cardiovasc. Imaging (Under review), (2023)

[57] Klein, A., García, M.: Diastology. Clinical Approach to Heart Failure with Preserved Ejection Fraction. Elsevier, Nederlands (2020)

[58] Fyrenius, A., Wigström, L., Ebbers, T., Karlsson, M., et al.: Three dimensional flow in the human left atrium. Heart 86(4), 448–455 (2001) https://doi.org/10.1136/heart.86.4.448

[59] Kilner, P.J., Yang, G.Z., Wilkes, A.J., Mohiaddin, R.H., et al.: Asymmetric redirection of flow through the heart. Nature 404(6779), 759–761 (2000) https://doi.org/10.1038/35008075

[60] Goette, A., Kalman, J.M., Aguinaga, L., Akar, J., et al.: EHRA/HRS/APHRS/SOLAECE expert consensus on atrial cardiomyopathies: definition, characterization, and clinical implication. Europace 18(10), 1455–1490 (2016) https://doi.org/10.1093/europace/euw161

[61] Khalili, E., Daversin-Catty, C., Olivares, A.L., Mill, J., et al.: On the importance of fundamental Computational Fluid Dynamics towards a robust and reliable model of left atrial flows: Is there more than meets the eye? Preprint at https://arxiv.org/abs/2302.01716 (2023)

[62] Esmaily-Moghadam, M., Hsia, T.Y., Marsden, A.L.: A non-discrete method for computation of residence time in fluid mechanics simulations. Phys Fluids 25(11), 110802 (1994) https://doi.org/10.1063/1.4819142

[63] Bolger, A.F., Heiberg, E., Karlsson, M., Wigström, L., et al.: Transit of blood flow through the human left ventricle mapped by cardiovascular magnetic resonance. J. Cardiovasc. Magn. Reson. 9(5), 741–747 (2007) https://doi.org/10.1080/10976640701544530

[64] Eriksson, J., Carlhäll, C.J., Dyverfeldt, P., Engvall, J., et al.: Semi-automatic quantification of 4D left ventricular blood flow. J. Cardiovasc. Magn. Reson. 12(9) (2010) https://doi.org/10.1186/1532-429X-12-9

[65] Suh, G.Y., Les, A.S., Tenforde, A.S., Shadden, S.C., et al.: Quantification of particle residence time in abdominal aortic aneurysms using magnetic resonance imaging and computational fluid dynamics. Ann. Biomed. Eng. 39(2), 864–883 (2011) https://doi.org/10.1007/s10439-010-0202-4

[66] Zhang, L.T., Gay, M.: Characterizing left atrial appendage functions in sinus rhythm and atrial fibrillation using computational models. J. Biomech. 41(11), 2515–2523 (2008) https://doi.org/10.1016/j.jbiomech.2008.05.012

[67] Zingaro, A., Dede, L., Menghini, F., Quarteroni, A.: Hemodynamics of the heart’s left atrium based on a variational multiscale-LES numerical method. Eur. J. Mech. B/Fluids. 89, 380–400 (2021) https://doi.org/10.1016/j.euromechflu.2021.06.014

[68] Jia, D., Jeon, B., Park, H.B., Chang, H.J., Zhang, L.T.: Image-based flow simulations of pre-and post-left atrial appendage closure in the left atrium. Cardiovasc. Eng. Technol. 10(2), 225–241 (2019) https://doi.org/10.1007/s13239-019-00412-7

[69] Grigoriadis, G.I., Sakellarios, A.I., Kosmidou, I., Naka, K.K., et al.: Wall shear stress alterations at left atrium and left atrial appendage employing abnormal blood velocity profiles. In: Conf. IEEE. Eng. Med. Biol. Soc., pp. 2565–2568 (2020). https://doi.org/10.1109/EMBC44109.2020.9175235

[70] Castañer, E., Gallardo, X., Rimola, J., Pallardó, Y., et al.: Congenital and acquired pulmonary artery anomalies in the adult: Radiologic overview. Radiographics 26(2), 349–371 (2006) https://doi.org/10.1148/rg.262055092

[71] Lyen, S., Wijesuriya, S., Ngan-Soo, E., Mathias, H., et al.: Anomalous pulmonary venous drainage: A pictorial essay with a CT focus. J. Congenit. Heart. Dis. 1(7) (2017) https://doi.org/10.1186/s40949-017-0008-4

[72] Mlczoch, J., Zutter, W., Keller, R., Herzog, H.: Influence of lung resection on pulmonary circulation and lung function at rest and on exercise. Respiration 32(6), 424–435 (1975) https://doi.org/10.1159/000193675

[73] D’Alessandro, N., Masci, A., Andalò, A., Dede, L., et al.: Simulation of the hemodynamic effects of the left atrial appendage occlusion in atrial fibrillation: Preliminary results. In: 2020 Computing in Cardiology, pp. 1–4 (2020). https://doi.org/10.22489/CinC.2020.302

[74] Cheng, C.P., Herfkens, R.J., Taylor, C.A., Feinstein, J.A.: Proximal pulmonary artery blood flow characteristics in healthy subjects measured in an upright posture using MRI: the effects of exercise and age. J. Magn. Reson. Imaging 21(6), 752–758 (2005) https://doi.org/10.1002/jmri.20333

